# A metabolic sum rule dictates bacterial response to short-chain fatty acid stress

**DOI:** 10.1101/2022.08.31.506075

**Authors:** Brian R. Taylor, Vadim Patsalo, Hiroyuki Okano, Yihui Shen, Zhongge Zhang, James R. Williamson, Joshua D. Rabinowitz, Terence Hwa

## Abstract

Short-chain fatty acids (SCFAs) such as acetate accumulate in fermentative environments, inhibiting many types of bacteria. While it is known that cells accumulate SCFAs to high concentrations internally, the cause of SCFA toxicity is not understood. By forcing *Escherichia coli* cells to accumulate a variety of “useless metabolites”, we establish via extensive ‘omic analysis a metabolic sum rule, by which the accumulation of exogenous metabolites such as acetate forces the depletion of endogenous metabolites. The latter leads to bottlenecks in biosynthesis, manifested as a simple linear relation between useless metabolite accumulation and growth reduction. Guided by quantitative models, we show that acetate-stressed cells optimize growth by partially acidifying their own cytoplasm, which reduces acetate accumulation, restoring the endogenous metabolites as allowed by the sum rule.

## INTRODUCTION

Short chain fatty acids (SCFAs) such as acetate accumulate in a variety of microbial environments. *E. coli* excretes acetate already during aerobic growth on glucose (*1*). If allowed to grow in conditions of excess carbon, the growth of *E. coli* becomes inhibited by the acetate it excretes. SCFAs are the end products of fermentative metabolism, which are found at inhibitory concentrations in environments such as the mammalian gut (*2–5*) and high-density biotechnology fermenters (*6, 7*). Additionally, SCFAs are used to inhibit the growth of microbes for food preservation (*8*). It is thus desirable to understand and modify bacterial tolerance to SCFAs. Particularly in biotechnology, the production of recombinant proteins by *E. coli* is limited by the acetic acid excreted (*6, 9, 10*). Furthermore, the industrial applications of high-value SCFAs produced by microbes is of significant interest, but the sensitivity of microbes to SCFAs remains a core problem in managing production costs (*11*).

The desire to understand the cause of SCFA stress has motivated several studies in *E. coli* with particular focus on acetate. Several identified causes include a drop of cytoplasmic pH (*12*), increase in cytosolic concentrations of SCFA (*13, 14*), and various metabolic effects (*14–16*). Yet, it is not known to what extent these causes quantitatively contribute to the growth rate reduction caused by the SCFAs. Nor is it known how *E. coli* combats these effects. In this study, we characterize these various causes quantitatively and integrate them into a mathematical model that reveals *E. coli*’s stress-resistance strategy and captures the growth rate behavior of acetate stressed cells.

## RESULTS

We characterized the toxic effect of medium acidity by growing *E. coli* K-12 cells in minimal glucose medium set to different pH using different buffers (see **Methods 2.1**). Growth was hardly affected down to pH 5, but dropped rapidly below that (purple symbols, **Fig. S1a**). However, in the presence of acetate, cells were more strongly affected by pH, with the effect being stronger for higher concentrations of acetate (brown and red symbols, **Fig. S1a**).

### SCFA-imposed growth defect depends exclusively on the concentration of the neutral acid

To explore the combined effect of acetate and pH further, we performed a systematic scan of these two variables. For each fixed medium pH (from 5 to 7), the growth rate steadily decreased as the medium acetate concentration was increased, with a stronger acetate sensitivity at lower pH (**Fig. 1a**). The combined effects of pH and acetate on growth rate can be collapsed onto a single curve if we plot growth rate against the concentration of acetic acid in the medium (**Fig. 1b**) where the acetic acid concentration, [HAc], is calculated from the Henderson-Hasselbalch equation (*17*),

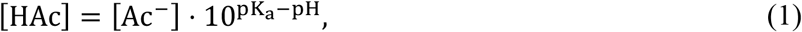

with pK_a_ ≈ 4.77 (*18*). The growth rate dependence on the acid form was shown previously for *E. coli* grown with benzoate (*19*).

**Figure 1.**
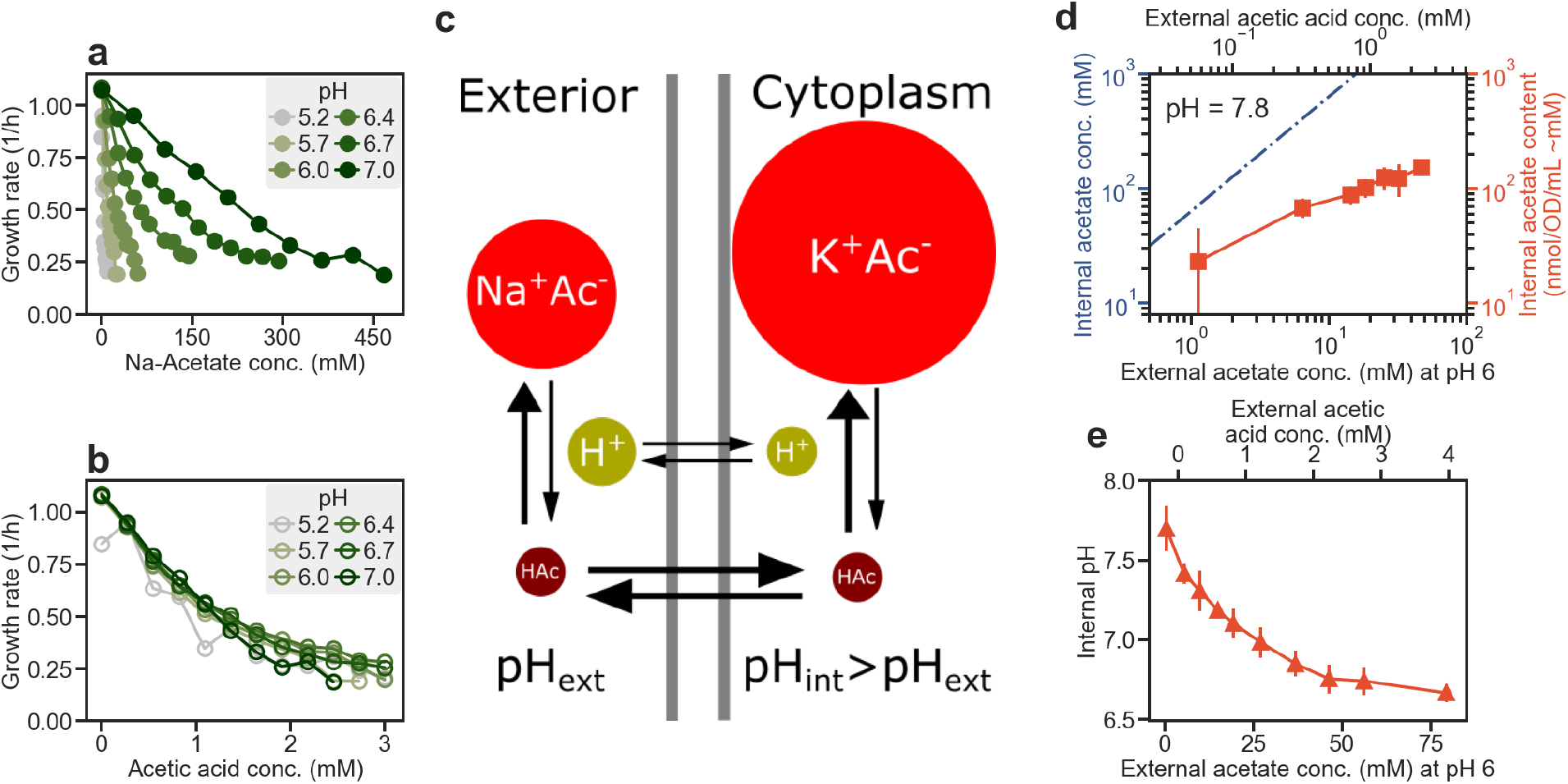
Effect of pH and acetate on bacterial growth. Unless otherwise indicated, *E. coli* K-12 NCM3722 cells were grown exponentially in glucose minimal media, at various pH fixed by phosphate buffer. Cultures were grown in either test tubes or Tecan microplate reader; see **Methods 2**. **a)** For several fixed medium pH, growth rate of the culture was determined for a range of added sodium acetate concentrations. (Excretion of acetate during growth, on the order of 1-2 mM (**Fig. S4h**), did not affect the medium pH.) **b)** The data in panel a is replotted against the concentration of acetic acid [HAc] using Eq. (1). **c)** Model of acetate equilibrium across the cell membrane adapted from Ref. (*13*). Acetate is in equilibrium with acetic acid on each side of the membrane according to Eq. (1). Acetic acid, due to its small size and lipophilicity, is membrane permeable. The cell controls its internal pH through transporters and the electron transport chain. A moderate difference between the internal and external pH results in a huge difference between the potassium-/sodium-acetate concentrations according to Eq. (2). **d)** Accumulation of internal acetate (right-axis) in NCM3722 cells; see **Fig. S2a,b** and **Methods 3.2** for details. The data is converted to internal acetate concentration (left y-axis) based on the measured water content (**Fig. S2e**). The dashed line indicates the internal acetate concentrations calculated according to Eq. (2) for internal pH fixed at 7.8, external pH = 6, and ΔpK*_a_* = 0. **e)** Internal pH was measured with the ratiometric reporter, pHluorin (26) for acetate stressed cells (HE616); see **Fig. S3a,b**. The results here also agreed well with the classical biochemical measurement using radiolabeled acid (**Fig. S3c**). Data in panels d and e are binned for similar x-axis values, with the bins containing data from n ≥ 3 biological replicates. Error bars are calculated from the standard deviation.

We next tested the toxic effect of several other weak organic acids. Similarly, to acetate, propionate and butyrate are two short-chain fatty acids (SCFA) that have very similar pK_a_ as acetate (*18*) and accumulate to high concentrations in the mammalian gut, reaching approximately 25 mM each compared to 60 mM for acetate (*20, 21*). Under a fixed pH, our cells responded similarly to butyrate as acetate, but exhibited somewhat increased sensitivity to propionate (blue and green circles, **Fig. S1b**). In comparison, the less commonly encountered benzoate, which also has similar pK_a_ (*18*), is much more toxic (purple, **Fig. S1b**). We will focus on acetate as the quintessential SCFA in this study.

Because high concentrations of SCFAs are commonly encountered in anaerobic conditions (*20, 22*), we also tested the acetate tolerance of anaerobically grown cells in glucose minimal medium at fixed pH of 6. Anaerobically grown cells exhibited similar growth rates as aerobically grown cells under acetate stress (**Fig. S1c**), suggesting that the cause of acetate toxicity is common to aerobiosis and anaerobiosis. We also characterized how a well-studied common gut bacterium, *Bacteroides thetaiotaomicron*, responded to acetate. We grew this obligate anaerobe in glucose medium with different acetate concentrations and pH. Like *E. coli*, the response collapsed to a single curve when plotted against acetic acid (**Fig. S1e**). Together, these results suggest that the toxicity to weak organic acids is a common phenomenon. The degree of toxicity is driven exclusively by the concentration of the neutral acid form, although the quantitative extent of the toxicity is dependent on the bacterial species and the identity of the acid.

### Acetate anions accumulate extensively in the cytoplasm with a modest drop of internal pH

We next turn to the consequence of elevated concentration of acetic acid, the source of the growth defect as established in **Fig. 1b**. Since acetic acid is a small, lipophilic molecule that can freely diffuse across lipid membranes (*23*) relatively quickly, the concentration of acetic acid is rapidly equilibrated inside and outside the cell. Given the relationship between acetic acid and acetate ions from Eq. (1), a simple relation was proposed (*13*) between the internal and external acetate concentrations (**Fig. 1c**):

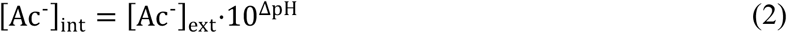

Here, ΔpH ≡ pH_int_ – pH_ext_ is the differences between the intra- and extra-cellular pH. Because the cell can control its internal pH through proton transport and the electron transport chain (*24*), a substantial difference between the internal and external pH can exist when the latter is low. As proposed by Russell (*13*), this pH difference would in turn result in substantial difference in the internal and external acetate concentration. Assuming that the internal pH is fixed at the unstressed value of ∼7.8 (see below), Eq. (2) predicts a huge build-up of internal acetate across the range of acetate stress applied in our experiments (dashed line, **Fig. 1d**), reaching ∼1 M at an external concentration of 20 mM.

We directly determined the internal acetate content for cells growing in this acetate range at medium pH=6 (**Fig. S2a,b**). The accumulated acetate content is shown as red squares in **Fig. 1d** (right y-axis) and charge is balanced by increased accumulation of potassium ions (**Fig. S2c,d**), suggesting that the accumulation of potassium is associated with the acetate accumulation, as a result of the acetic acid being neutralized. We additionally measured the cytoplasmic water content using radiolabeling (**Fig. S2e**), finding it to remain at ∼1 μL/OD/mL (∼2x cell dry weight (*25*)), largely independent of the degree of acetate stress; thus, the measured acetate content (in nmol/OD/mL) converts simply to mM concentration (**Fig. 1d**, left y-axis). The accumulated acetate was much lower than that expected if cells had maintained their internal pH at 7.8 (compare red squares and dashed line, **Fig. 1d**). Also, internal acetate scaled sub-linearly with the external acetate concentration, indicating that the internal pH and/or ΔpKa were changing under different degrees of acetate stress as reported in early studies (*13, 14*). We confirmed a drop in internal pH for these acetate-stressed cells (red triangles, **Fig. 1e**) using a pH-sensitive ratiometric GFP, pHluorin (*26*) (**Fig. S3a,b**), cross-validated with the classical radioactivity method (*27, 28*) (**Fig. S3c**). Given measurements of both the internal pH and acetate concentration, we calculated the internal pK_a_ using Eq. (1) and found a 0.1-0.2 unit shift from its accepted value across the range of extracellular acetate (**Fig. S3d**).

### Rapid and slow adaptation of the metabolome to sudden acetate stress

We next turn to the effect of acetate accumulation on the metabolome. It was previously reported (*14*) that acetate accumulation decreased the level of internal glutamate. We followed the cellular response kinetically after 30 mM of sodium acetate was spiked into exponentially growing cultures (with medium pH set to 6, which did not affect growth in the absence of acetate; see **Fig. S1a**). After a very brief transient of about 15 minutes, the instantaneous growth rate reached ∼0.3/h (**Fig. 2a** blue circles), close to the steady-state growth rate in the condition with acetate (horizontal dashed line).

**Figure 2.**
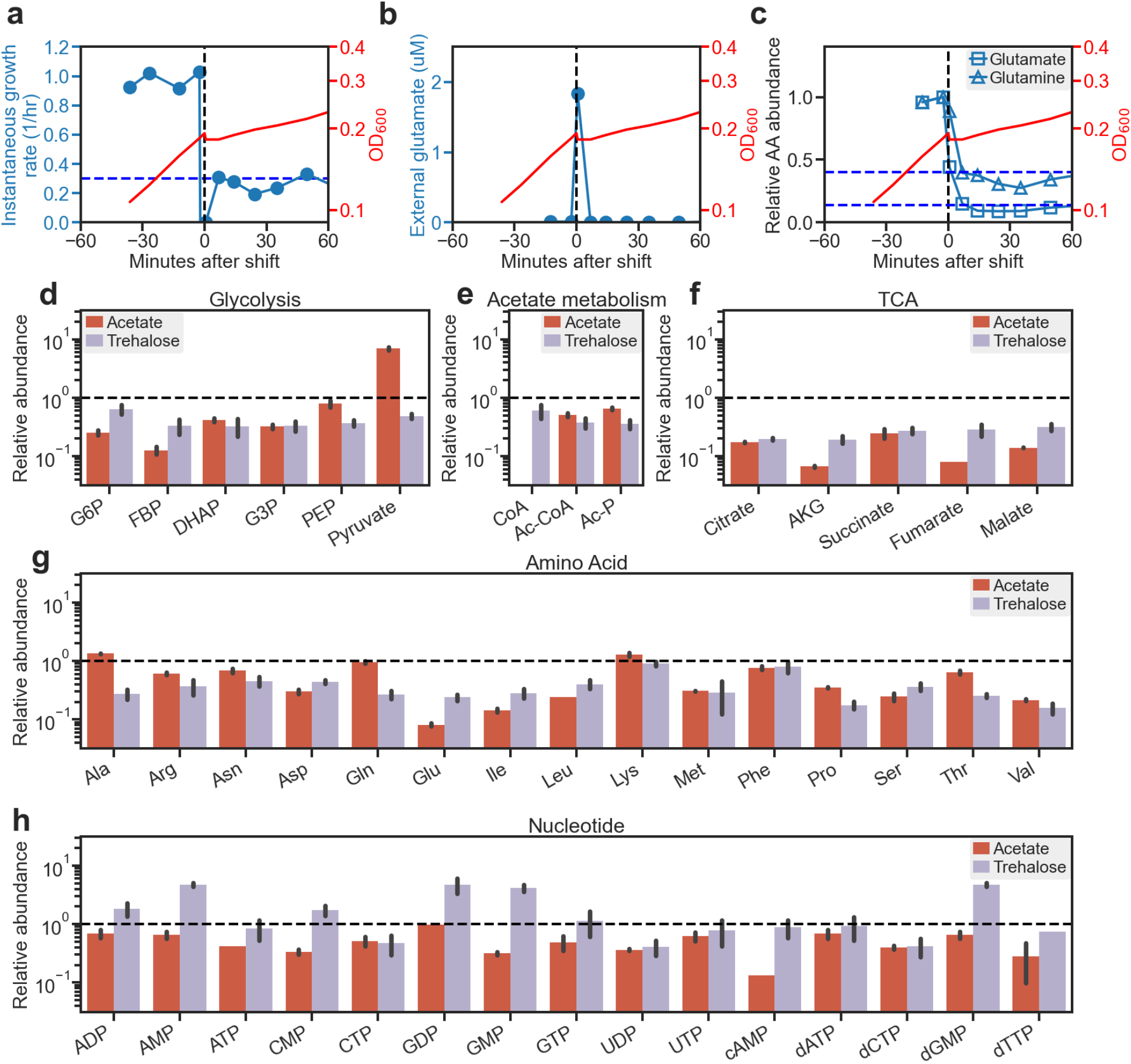
Transient and steady-state changes in metabolite concentrations. **a-c)** NCM3722 cells growing in steady state in glucose minimal medium at pH 6 without acetate were exposed to 30 mM acetate at time 0 and followed thereafter. **a)** Instantaneous growth rate was calculated as the centered logarithmic derivative of OD_600_ with respect to time. **b)** External glutamate pool was measured by HPLC in filtered media. **c)** Relative internal abundances of glutamate and glutamine were measured for cells by HPLC (**Methods 3.4)**. **d-h)** Red bars: NCM3722 cells growing in steady-state were assayed for metabolite concentrations in pre-shift and post-shift acetate conditions; growth rates were 0.93/h and 0.30/h, respectively. [Note that CoA was not detected under acetate stress; this was not due to poor detection limit as the pools of both CoA and its precursor Panthothenic acid dropped sharply in medium with lower amount of sodium acetate added, becoming undetectable above 15 mM (**Table S3a**).] Purple bars: HE647 or HE650 cells growing in glucose medium at pH 6 and no acetate, supplemented by zero and 50 ng/mL of the inducer cTc which controls internal trehalose accumulation; see **Fig. S9**. Metabolites were quantified by LC-MS following fast filtration (*30*). For the relative fold-change, a ^13^C reference was mixed in with the collected samples; see **Methods 4.1**. The metabolomic data are given in **Table S3a** for acetate-stressed cells and **Table S3c** for cells with trehalose overdose. Error bars are calculated from the standard deviation.

The rapid adaptation of cell growth to the sudden acetate stress was mirrored by the rapid response of some of the metabolites measured by HPLC (**Methods 3.3**). Immediately after acetate addition, a small amount of glutamate was excreted to the medium, but then disappeared within 4 minutes, likely taken back by the cells (blue circles, **Fig. 2b**). Sudden and substantial decreases were also observed in internal glutamate and glutamine pools immediately following acetate addition. Glutamate, the most abundant metabolite in growing cells (*30*) and used as a substrate for the synthesis of most other amino acids (*31*), dropped to under 10% of its steady state concentration within four minutes of the acetate addition (**Fig. 2c** squares), and remained there into the steady state (dashed horizontal line). Similarly, glutamine, also used in amino acid synthesis, dropped ∼30% to its post-acetate steady state concentration within a similar timescale (**Fig. 2c** triangles and dotted line). The immediate drop of these two key amino acids likely resulted from some bottleneck in their synthesis caused by acetate addition (see below).

The addition of acetate also led to a large disruption in carbon utilization, which unlike the quick changes in growth rate and glutamate/glutamine pools (**Fig. 2a-c**), remained perturbed for an extended period before stabilizing to their steady-state values. This involved a much-increased specific glucose uptake (i.e., a reduced glucose yield) for several hours following acetate addition (blue symbols, **Fig. S4a-b**), compared to that of either the pre- or post-acetate steady state (green and brown symbols, **Fig. S4b**). The glucose influx, which is the product of growth rate and specific uptake, remained over 2x above the level in the post-acetate steady state during the transient phase (**Fig. S4c**). The excess specific glucose uptake coincides with the severe excretion of pyruvate (**Fig. S4d**), likely resulting from the buildup of internal pyruvate due to the depletion of CoA by the large amount of acetate entering the cytoplasm (**Fig. S5a**), and an incomplete inhibition of glucose transport by the PTS system (**Fig. S5b,c**). The buildup of pyruvate disappeared in the steady state with acetate, where pyruvate excretion was much reduced (brown symbols, **Fig. S4f**), and the glucose yield was similar to that before acetate addition (brown symbols, **Fig. S4b**). Furthermore, adaptation to acetate also leads to a slow reversal from acetate excretion to acetate uptake (**Fig. S4g-i**). The cause of this slow adaptation likely involves the dilution of several catabolic proteins to bring down the glucose influx. This scenario is discussed in detail in **Fig. S5,S6**.

We sought to cross-check the above scenario by performing metabolomic experiments, characterizing the relative abundance of 119 intracellular metabolites in the pre- and post-acetate steady state conditions using LC-MS (*30, 32*) (see **Table S3a** and **Methods 4**). We found at least several-fold decreases in the concentrations of many metabolites in glycolysis and the TCA cycle (red bars, **Fig. 2d-f**), particularly with CoA dropping to undetectable level in the presence of acetate. The one exception is pyruvate which increased in concentration by nearly 10-fold, reflecting the observed pyruvate excretion (**Fig. S4d**). The concentrations of 13 out of 15 measured amino acids decreased as well, with only alanine and lysine increasing slightly (red bars, **Fig. 2g**). Furthermore, the concentrations of most nucleotides decreased including cAMP, the indicator for global carbon status (*33*) (red bars, **Fig. 2h**). The latter is expected based on the increase in pyruvate (**Fig. S4d**) and the known inhibition of cAMP synthesis by keto-acids including pyruvate (*35*). Thus, acetate-stressed cells perceived themselves in a glucose-replete state.

The data on the changes in metabolite concentrations additionally allowed us to formulate a hypothesis on how the excretion of glutamate immediately following acetate addition could be stopped in a few minutes (**Fig. 2b,c**), in stark contrast to pyruvate and alanine excretion which lasted for several hours (**Fig. S4d,eS5**). As elaborated in **Fig. S6**, glutamate is supplied by the anaplerotic flux catalyzed by Phosphoenolpyruvate Carboxylase (Ppc). The latter is strongly activated allosterically by fructofuranose-1,6-biphosphate (fbp) (*36, 37*), a key glycolytic intermediate which is established to sense glycolytic flux (*38*). Thus, a drop in the rate of glucose uptake (**Fig. S4c**) is sufficient to reduce the fbp pool and consequently limit the anaplerotic flux that fuels the biosynthesis of 10 amino acids including glutamate derived from TCA intermediates (**Fig S6**).

### The proteome of acetate-stressed cells corroborates bottleneck in glutamate synthesis

The immediate drop in the internal glutamate and glutamine pools following acetate addition (**Fig. 2c**) concomitantly with the immediate drop in the instantaneous growth rate (**Fig. 2a**), both to their post-acetate steady state levels, suggest that the reduced pools of these and possibly other amino acids could be responsible for the reduction in cell growth. This could result directly from the reduced charging levels of the tRNA for glutamate or glutamine, indirectly through the effect of glutamate and/or glutamine on the synthesis of other amino acids through trans-amination and trans-amidation reactions (*31*), through glutamate’s role in internal pH regulation (*14*), or through other global effects as glutamate is the major anion in the cytoplasm (*39*).

To see whether the reduced glutamate pool could drive the observed growth reduction, we recall previous studies of *E. coli* subjected to synthetic internal limitation in glutamate biosynthesis (*29, 35, 40*). Like the acetate-stressed cells, cells with limited glutamate synthesis exhibited reduced pools of amino acids including glutamate itself, and reduced cAMP level (*40*). To further investigate possible similarities between cellular responses to these two modes of growth limitations, we characterized the proteome of acetate-stressed cells using mass spectrometry (Ref. (*29*), **Method 6**) and compared the results to those previously reported for limited glutamate synthesis, as well as to those for limited sugar uptake and limited protein synthesis as controls (*29*).

We first compared the pattern of fold-changes in protein abundances (**Fig. S7a-c, Table S2a**). Those fold-changes for acetate-stressed cells were most correlated with the fold changes for cells limited by glutamate synthesis (**Fig. S7e**). This similarity of protein expression pattern to limited glutamate synthesis is also observed when proteins are grouped according to physiological function (orange symbols in **figS. S7f, S8**). In particular, amino acid biosynthesis, TCA and glyoxylate shunt were the functional groups that increased the most for both acetate stress and glutamate synthesis limitation while motility and ribosomal proteins were decreased. Altogether, these results suggest that acetate exerts its toxic effect broadly on biosynthesis, in ways similar to that obtained by a limiting glutamate synthesis.

### Quantitative global effects of acetate stress on the metabolome

We next investigated the link between glutamate depletion and acetate stress, which occurred immediately following acetate addition (**Fig. 2c**). To do so, we quantified the absolute concentrations of 72 metabolites which are a part of *E. coli* central carbon metabolism and amino acid biosynthesis (see **Table S3a),** based on the measurements of Ref. (*32*) (**Methods 4**). We found that their total concentration dropped from 350 mM to 100 mM, amounting to a total decrease of 250 mM after including the neutralizing potassium ions associated with the large amounts of glutamate and aspartate (grey and black bar, **Fig. 3a**). The total decrease was comparable to the total amount of potassium acetate accumulated in the cytoplasm (red and pink bars in **Fig. 3a**).

**Figure 3.**
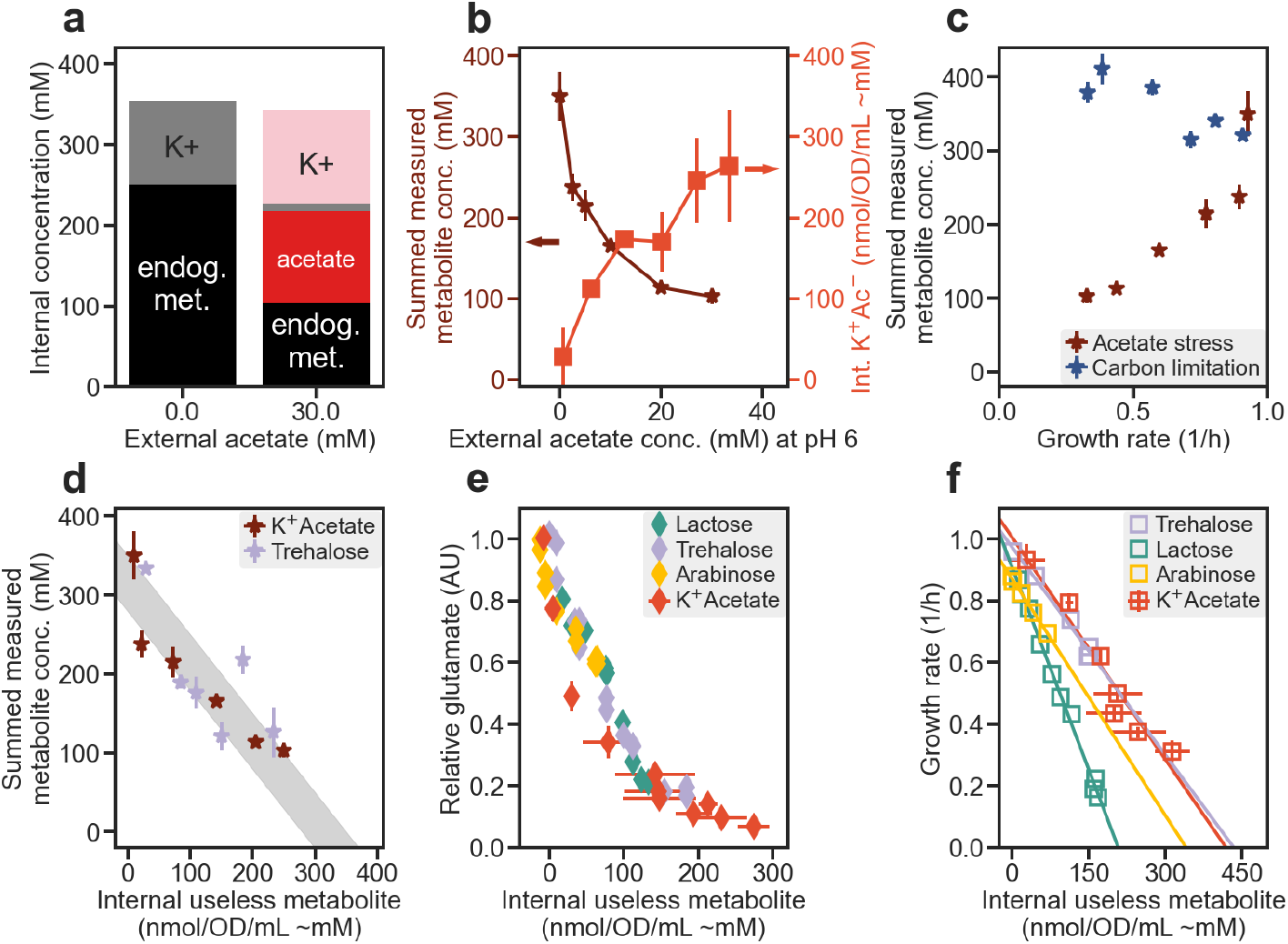
Effect of useless metabolites. **a)** For NCM3722 cells grown in glucose minimal medium at pH 6, the absolute concentrations of 72 internal metabolites determined by LC-MS measurements (**Methods 4**) were summed and represented as the height of the black bars for medium without and with 30 mM of sodium acetate. The concentrations of the neutralizing potassium ions for glutamate and aspartate are represented as the height of the grey bars. The concentration of the acetate accumulated, measured in **Fig. 1d**, is shown as the red bar, and its neutralizing potassium ion concentration is shown as the pink bar. **b)** Same as panel a, but for different amounts of external acetate. The sum of the abundances of 72 internal metabolites, together with the neutralizing potassium ions for glutamate and aspartate but not including potassium acetate, are shown as brown stars (left y-axis). Data for the internal potassium acetate (red squares, right y-axis) are taken to be twice of the measured internal acetate shown in **Fig. 1d**. **c)** The sum of the measured metabolites under acetate stress (brown stars, same as in panel b) and under carbon-limited growth (blue stars), plotted against the respective growth rates. Carbon limitation was implemented by titrating glucose uptake (*1*) for cells grown in glucose minimal media (see **Methods 2.1**, with data provided in **Table S3b**). The cellular water content under these conditions are similar (**Fig. S2e,f**). **d)** Sum of the abundances of all metabolites determined by metabolomics for trehalose overdose (purple stars, again including the neutralizing potassium ions for glutamate and aspartate) is plotted against the total trehalose content for strains HE647/HE650 (**Fig. S9)**. Grey band represents the range bounded by a fixed metabolite concentration ranging from 300-350 mM. The data are provided in **Table S3c**. The data for acetate stress (brown stars) is the same as that shown in panel b and c but plotted here against potassium acetate (red squares in panel b). **e)** Effect of useless metabolites on internal glutamate, due to acetate stress (red diamonds), trehalose overdose (purple diamonds), lactose overdose (teal diamonds), and arabinose overdose (orange diamonds), in strains NCM3722, HE647/HE650, HE620, and HE639, respectively; see **Fig. S9** and **Table S1a** for description. Useless metabolites on x-axis are the measured amounts of potassium acetate, lactose, trehalose, and arabinose accumulated in the respective strains. The measured acetate amount is multiplied by two to include the additional increase of potassium cations needed to neutralize the acetate anions (see **Fig. S2d**). **f)** Same as panel e but showing the corresponding growth rates. All growth was performed in phosphate-buffered media at pH 6. Data in panels b, e and f are binned for similar x-axis values with the bins containing data from n ≥ 3 biological replicates. Data in panels c and d are grouped according to the same applied stress level and each point represents n = 2 biological replicates. Error bars are calculated from the standard deviation.

To see how the amount of endogenous metabolite loss depended on the degree of acetate stress, we repeated the LC-MS measurements for cells growing in steady state with different levels of sodium acetate in the medium at pH 6. The results are listed in **Table S3a**, with the total measured metabolites (including the potassium counter ions) plotted as the brown stars in **Fig. 3b**. The total concentration of measured metabolites decreased gradually as the concentrations of internal potassium acetate accumulated (red squares, **Fig. 3b**). We further characterized the same metabolites in cells limited in sugar uptake, which have similar water content (**Fig. S2f**). Comparing the total concentrations of the measured metabolites in these two cases at the same growth rate (**Fig. 3c**, **Table S3a,S3b**), the drop of metabolites in acetate-stressed cells (brown stars) is conspicuous.

The global reduction of such a significant fraction of endogenous metabolites under acetate stress can be rationalized by the hypothesis that the total internal metabolite concentration is maintained at an approximately constant level in our experimental conditions which has external osmolarity in the range of 280-350mM, the range known for healthy mammalian gut (*41*). The maintenance of total internal metabolite concentration may result from the cells reaching an upper limit in turgor pressure, which sets the difference between the internal and external osmolarity (*42*). Indeed, plotting the total concentration of the measured metabolites against the accumulated potassium acetate concentration (brown stars, **Fig. 3d**) yields a relation consistent with the total concentration being maintained at 300-350 mM (grey band). Consequently, when the cytoplasm is forced to accommodate large amounts of potassium acetate, a “useless metabolite” for *E. coli* from the metabolic perspective, the concentrations of normal metabolites must decrease. This decrease would provide a possible cause for growth defect under acetate stress: because many metabolite concentrations are poised near the *K_m_* of their respective enzymes in unstressed growth conditions (*30*), a reduction of normal metabolite pools under acetate stress would reduce the metabolic flux catalyzed by these enzymes and thereby slow down growth. This mechanism of growth reduction would account for the immediate drop of growth rate after the addition of acetate (**Fig. 2a**), concomitant with the immediate drop of the two internal concentrations of amino acids measured, glutamate and glutamine (**Fig. 2b,c**). This mechanism could also readily account for the upregulation of biosynthetic enzymes found in the proteome analysis (**figS. S7, S8**), which could result from end-product feedback control, either directly due to reduction of intermediate metabolite pools or indirectly through reduced trans-amination arising from the reduced glutamate pool.

### Generic effects of useless metabolites on the metabolome and cell growth

To test this “useless metabolite” hypothesis directly, we constructed strains to accumulate three distinct metabolites, trehalose, lactose and arabinose, respectively shown in **Fig. S9a-c** (see also **Methods 1)**. These molecules are “useless” to the cells because the strains have been designed to accumulate but not metabolize them (**Table S1a**). Additionally, these molecules are relatively inactive; in particular, trehalose is a molecule that accumulates to high concentrations in osmotically stressed cells (*43*). We measured the internal lactose, arabinose, and trehalose content accumulated (**Methods 3.2**) and established that the accumulation of these metabolites can indeed be titrated across a broad range, with a concomitant decrease in the glutamate pool (**Fig. S9d-f**). Moreover, the growth rate dropped as well, while it changed little for a control where GFP is expressed by the same system (**Fig. S9g-i**). We found the accompanying cytoplasmic water content to change little (**Fig. S2g**), so that the intracellular content measured (nmol/OD/mL) converts directly to concentration (mM) as with acetate stress. Notably, growth inhibition due to the overdose of lactose (**Table S2b**) resulted in a proteomic phenotype similar to that of acetate stress (**Fig. S7e,g**).

We characterized the metabolome of cells subjected to various degrees of trehalose overdose by LC-MS, and obtained the absolute abundance of the 72 metabolites as mentioned above (**Table S3c**, **Methods 4.1**). Their abundances relative to unstressed cells are shown as purple bars in **Fig. 2d-h** for the ease of comparison to those in acetate-stress cells. As in the response to acetate stress, many metabolite pools were reduced for cells under trehalose overdose, even for pyruvate and alanine, whose pools increased under acetate stress. Surprisingly, some of the nucleotide pools increased (**Fig. 2h**). However, they hardly affected the energy charge (*34*) (**Fig. S3i**), reflecting that these cells, like acetate-stressed cells, do not have an “energy problem”.

The total concentration of measured metabolites (including potassium counter ions for glutamate and aspartate) for trehalose overdosed cells are shown as purple symbols in **Fig. 3d**. As predicted by the “useless metabolite” hypothesis, the total concentration decreased as the concentration of trehalose, the useless metabolite, increased. Strikingly, the amount of decrease is similar between trehalose overdose and acetate-stress (**Fig. 3d** brown symbols). We also measured glutamate using HPLC for the lactose and arabinose overdoses and found that, similarly to trehalose overdose and acetate stress, the concentration of glutamate decreased as lactose or arabinose accumulated, in similar ways to the effect of trehalose and potassium acetate (**Fig. 3e).** Thus, regardless of the identity of the useless metabolite, its accumulation quantitatively dictates the decrease of internal metabolites, in accordance with the useless metabolite hypothesis.

We next made similar plots for the measured growth rate against the internal concentration of each useless metabolite (**Fig. 3f**). A striking linear decrease of growth rate (*λ*) is obtained in each case, of the form

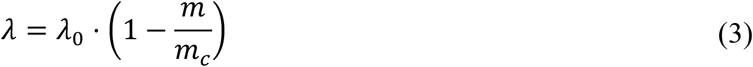

where *m* is the internal concentration of the useless metabolite, *m_c_* is the inhibitory concentration where growth rate is extrapolated to vanish, and *λ*_0_ is the stress-free growth rate. The inhibitory concentrations are surprisingly different for the different substances: accumulation of lactose led to 2x steeper reduction in growth compared to trehalose even though both are neutral disaccharides while the accumulation of arabinose led to a growth rate reduction in between the other two. Since the effect of these useless metabolites on the internal metabolites are similar (**Fig. 3e**), the results indicate that growth reduction by useless metabolites is not a sole consequence of their reduction of the normal metabolite pools. It is known that for metabolites accumulated to such high concentrations (e.g., 100 mM of disaccharides corresponds to ∼3.4% w/v), general properties of the cytoplasm may be affected, including, e.g., the solvation and hence the activity of proteins (*44*), which we refer to collectively as the “solvent effect”. Trehalose and potassium acetate could be metabolites that the proteome of *E. coli* is well-adapted to, as an osmolyte for trehalose (*43*), and due to frequent exposure in the environment for acetate and other SCFAs (*19, 20*). The proteome may be less adapted to lactose or arabinose as high internal concentration of lactose or arabinose is not commonly encountered, thus resulting in lower inhibitory concentration.

### Tradeoff between useless metabolite accumulation and internal pH drop

The fact that acetate accumulation resulted in linear growth reduction similar to the accumulation of trehalose, arabinose, and lactose (**Fig. 3f**) is somewhat surprising given the fact that the accumulation of acetate in the cytoplasm can be adjusted by changing the internal pH; see Eq. (2). In principle, cells can simply set the internal pH to the same value as the external pH and thereby eliminate potassium acetate accumulation altogether. Here, we hypothesize that reducing the internal pH would cause a growth defect of its own, and *E. coli* actively seeks a compromise, reducing its internal pH partially (as seen in **Fig. 1e**) in response to external acetate, to reduce potassium acetate accumulation despite the cost of reduced internal pH itself. Specifically, we hypothesize that the internal pH is set in a way that the combined growth defect due to potassium acetate accumulation and internal pH drop is minimized (**Fig. 4b**).

**Figure 4.**
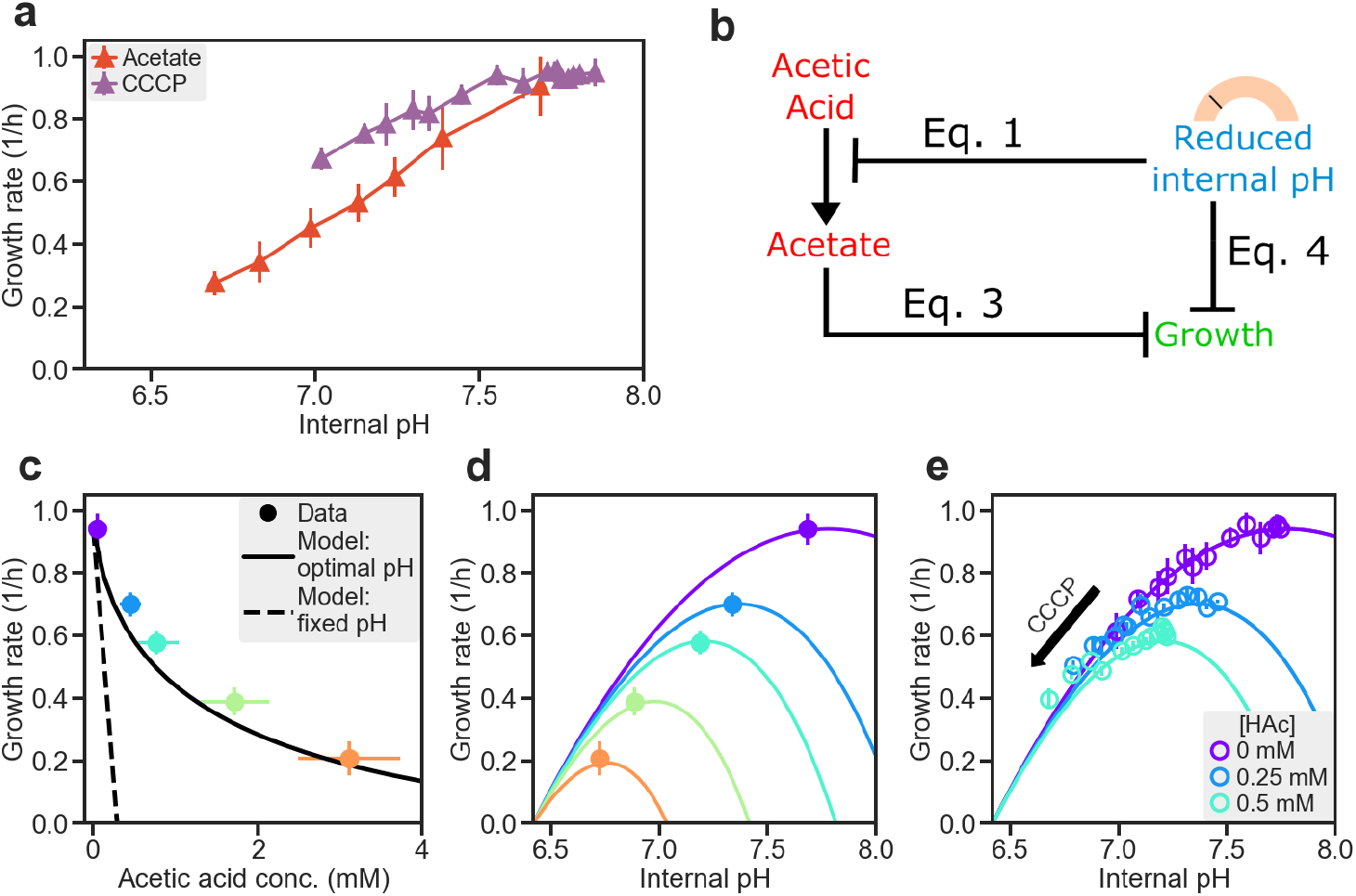
Model of acetate-pH tradeoff. **a)** The internal pH data for cells grown in glucose medium at pH 6 with various external acetate concentrations (data shown in **Fig. 1e**) is plotted against the growth rate of the corresponding culture (red triangles). Additionally, internal pH was measured for cultures of the same cells grown in varying amounts of CCCP (purple triangles) in the same phosphate buffer set to pH 6; see **Fig. S3f** and **Methods 5.1**. HE616 cells harboring pHluorin was used in order to collect the internal pH data. **b)** Schematic of the acetate-pH tradeoff model. **c)** The model output (solid line) quantitatively captures the effect of growth reduction by acetic acid (colored circles) with no adjustable parameter; see **Fig. S10** and **Supplementary Note** for details. Dashed line shows the expected growth rate if internal pH is fixed at an unstressed value of 7.78. **d)** Circles show the measured growth rate and internal pH for cultures in media with various acetic acid concentrations (same as **Fig. S1f**). Color indicates the acetic acid concentrations shown in panel e. Solid lines show the expected growth rate if the internal pH is set to the values indicated by the x-axis. **e)** The lines and the filled circles are the same as those shown in panel d. Open circles are the results of additional experiments with combined acetate and CCCP stresses. HE616 cells harboring pHluorin were grown in glucose media buffered to pH 6, with various concentrations of sodium acetate (indicated by the colors), each with a range of CCCP concentrations to perturb the internal pH. Data in panels b, c, d, and e are grouped according to the same applied stress level and contain n ≥ 3 biological replicates. Error bars are calculated from the standard deviation.

To test our hypothesis quantitatively, we first characterized the effect of reducing the internal pH on growth without involving useless metabolites. This was done by growing cells in glucose medium at normal pH in the absence of acetate, with the supplement of varying amounts of an uncoupler, Carbonyl cyanide m-chlorophenyl hydrazone (CCCP), which is known to reduce the internal pH (*45*). We found the growth rate to decrease along with the internal pH for increasing dose of CCCP (**Fig. 4a**). However, growth reduction due to CCCP was always smaller than that due to acetate stress at the same internal pH, consistent with our hypothesis that acetate toxicity resulted from a combination of a drop of internal pH and cytosolic accumulation of acetate.

We next developed a mathematical model incorporating the deleterious effect of accumulating potassium acetate as a useless metabolite, described by Eq. (3) (with *m* being the internal potassium acetate concentration), and with potassium acetate accumulation affected by the internal pH via Eq. (1). The deleterious effect of internal pH, pH_int_, alone on growth (**Fig. 4a**) is modeled by a parabolic dependence,

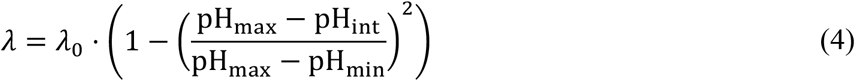

with the maximum set to pH_max_ = 7.78, and minimum at pH_min_ = 6.42, obtained as best-fit to data (**Fig. S10a**).

The joint effect of potassium acetate accumulation and internal pH reduction is assumed to be a product of the individual effects described by Eqs. (3) and (4) with the internal metabolite concentration *m* in Eq. (3) being potassium acetate, set as 2 × [*Ac*^−^]*_int_*, the latter given by Eq. (2) through the acetic acid concentration set by the external environment. The internal pH is controlled by the cell through proton import and export. The inhibitory concentration *m_c_* for potassium acetate as a useless metabolite is the lone unknown model parameter.

As described in **Supplementary Note**, this model can be solved analytically. The model predicts that for a substantial range of acetate stress, the growth rate and internal potassium acetate concentration are approximately linearly related, with an apparent inhibitory concentration,

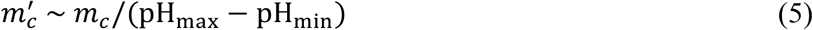

when the growth rate under acetate stress is extrapolated to zero. A one-parameter fit of the model to the acetate/growth-rate data fixed the inhibitory concentration to *m*^<^ ≈ 400 *mM* for potassium acetate. With this parameter, the model quantitatively captured the relationship between potassium acetate, pH, and growth rate, as shown in **Fig. S10b-d**. **Fig. 4c** shows the comparison of the predicted growth-rate dependence on acetic acid concentration, with the dashed line indicating the expected growth defect if internal pH was not reduced. Further, **Fig. 4d** shows for a number of acetic acid concentrations (different colors) what the growth rate would have been if internal pH were fixed at various levels (x-axis). Strikingly, the observed data lie closely to the growth rate maximum at each acetic acid level (circles of corresponding colors), self-consistently justifying the optimality assumption. To further test the model, we applied a “double stress”, by varying the amounts of CCCP to cells growing in glucose at different fixed acetic acid concentrations. **Fig. 4e** shows that the growth rate quantitatively followed the parabolic trajectories as predicted by the model, without invoking any new parameters. Together, these data establish that *E. coli* cells indeed set the internal pH to optimize the tradeoff between acetate accumulation and pH reduction.

## DISCUSSION

In this study, we systematically characterized the physiological, proteomic, and metabolic responses of the model bacterium *E. coli* to short-chain fatty acid (SCFA) stress. Extending previous results on benzoic acid (*19*), we established that the toxicity that acetate exerts is through the concentration of the neutral acetic acid, which is itself determined by the pH and the amount of acetate anion in the medium (**Fig. 1a,b**). Quantitatively similar effects are seen for other common SCFAs such as propionate and butyrate, and for aerobiosis and anaerobiosis (**Fig. S1**). Focusing on the effect of acetate, we find acetate anions accumulate intracellularly to hundreds of mM (**Fig. 1d**), accompanied by similar increase in the neutralizing potassium ions (**Fig. S2c**), along with a decrease in cytosolic pH (**Fig. 1e**).

Our analysis showed that acetate accumulation causes a defect in glutamate synthesis that leads to a depletion in cellular resources downstream. When cells are suddenly exposed to acetate at low pH, their glutamate and glutamine concentrations quickly drop within minutes of the exposure, concomitantly with a quick drop in growth rate (**Fig. 2a-c**). Additionally, many metabolites are depleted substantially, including all TCA intermediates and most amino acids detected (**Fig. 2d-h**). Glutamate, and to a lesser extent glutamine, is known to play a central role in the amino acid biosynthesis, taking part in the *de novo* biosynthesis of other amino acids through trans-amination and trans-amidation reactions (*31*). Previous studies have characterized the proteome and metabolome of cells whose growth were limited by glutamate synthesis (*29, 35*). The proteome of acetate-stressed cells resembles the proteomes of cells limited in glutamate synthesis (**Fig. S7, S8**). Their metabolomes also resemble each other, with many metabolites including glutamate decreasing as growth slows down. The biosynthetic defect does not appear to cause energy problems as both the carbon yield (**Fig. S4b**) and the energy charge (**Fig. S3i**) are maintained. Sufficient supply of carbon is signaled by a sharp drop in the level of cAMP, the indicator of carbon status (**Fig. 2h**).

Our quantitative metabolomic data led to the “useless metabolite” hypothesis as an explanation of the general metabolite depletion observed. Cells that accumulate potassium acetate experience an equivalent decrease in endogenous metabolites such that their total concentration remains constant at 300∼350 mM (**Fig. 3d**), similar to the osmolarity of the healthy mammalian gut which *E. coli* is native to (*41*). This observation was replicated for cells designed to accumulate synthetic “useless metabolites”, which excluded endogenous metabolites, ultimately leading to growth defects (**Fig. 3d,f** and Eq. (3)).

As the cell can control the amount of acetate accumulation through the internal pH, we hypothesize that the final effect of acetate on cell growth is obtained from a strategy that trades off two opposing effects of lowering internal pH: a direct deleterious effect on growth and a beneficial effect of reducing acetate accumulation. A simple mathematical model combining the effect of useless metabolites on reduced internal pH on cell growth is able to account for not only all of the existing data but also combined effect of acetate and internal pH changes without using free fitting parameters (**Fig. 4**).

*E. coli*’s strategy to partially lower its internal pH to mitigate acetate stress can be readily generalized to describe bacterial response to other permeable weak acids (**Fig. S1b**), whose intracellular concentrations are determined by the internal pH via Eq. (2). Our work indicates that the inhibitory effect of these weak acids, given by Eq. (5), is determined by a compromise of two deleterious factors: the inhibitory concentration (*m_c_*) of the acid as a useless metabolite, and the minimal internal pH (pH_min_) that the cell can tolerate. More detailed calculation for the half-inhibitory acetic acid concentration (**Fig. S10e,f** and **Supp Note**) predicts additional weak dependence on the value of the normal pH (pH_max_). The toxicity of the different weak acids presumably arises from their different disruptive effects as useless metabolites (e.g., via the afore-mentioned solvent effect), with *E. coli* adapting better to the SCFAs commonly encountered in the gut, compared to, say, benzoic acid, analogous to the differences in *m_c_* among trehalose, arabinose, and lactose (**Fig. 3f**). The similarity of the *m_c_* values under trehalose and acetate stresses to the osmolarity of the medium, which was set close to the healthy gut (*41*), suggests that metabolite concentrations are constrained by the environmental osmolarity, perhaps due to a limit on turgor pressure.

It is noteworthy that the cellular parameters (*m_c_*, *pH_min_*, and *pH_max_*) determining the toxicity of a weak acid are *global*, in the sense that they are determined by properties of numerous cytosolic proteins. The modification of these parameters would require changing the properties of many of those proteins and cannot be mitigated by expressing a few genes as in the case of osmolarity stress or low pH stress (*46*). Accordingly, adaptation to weak acid stress by e.g., lowering the minimal internal pH, would require such global modifications and be costly to cells grown in neutral pH. This fundamental tradeoff may underlie a bifurcation of bacterial responses in communities frequently exposed to weak acids stress: those which grow fast in normal pH but is sensitive to low pH, and others which grow slower in normal pH but has improved tolerance at lower pH (*2, 4*). Understanding these tradeoffs will be instrumental to managing and manipulating these microbes in environments ranging from the gut to bioreactors where fermentation products dominate (*5, 47–51*).

## Acknowledgements

We are grateful to helpful discussions with many colleagues during the course of this work, including Kapil Amarnath, Jonas Cremer, Adam Feist, Harry Flint, Matteo Mori, Jun Park, Bert Poolman, Jack Reddan, Amir Zarrinpar, Karsten Zengler, and the late George MacFarlane. We thank Hui (Tony) Sheng for early work on metabolomics, and Joan Slonczewski for providing JLS1105. This work is supported by the NIH through grant R01GM109069 (TH) and R35-GM136412 (JRW). JDR and YS acknowledge the support of the Ludwig Cancer Research.

## Author contribution

Conceptualization: BRT, VP, TH

Data curation: BRT, VP, YS

Formal analysis: BRT, VP, YS

Funding acquisition: JRW, JDR, TH

Investigation: BRT, VP, HO, YS

Model development: BRT, TH

Methodology: VP (proteomics), HO (measurements involving radioactivity, potassium measurements), ZZ (strain construction), YS (metabolomics), BRT (HPLC, fluorescence measurements, anaerobic cultures)

Project administration: TH

Supervision: TH

Visualization: BRT, TH

Writing – original draft: BRT, TH

Writing – review & editing: BRT, HO, YS, ZZ, JDR, TH

## Data and materials availability

Raw mass spectral data is deposited to massIVE and accessible with the accession code MSV000087288 (ftp://MSV000087288@massive.ucsd.edu, password aceticAcid), and will be made publicly accessible upon publication. All other data are available in the main text or the supplementary materials.

The metabolomics data is deposited in Metabolights under study number MTBLS5374 (www.ebi.ac.uk/metabolights/MTBLS5374)

## Supplementary Information

**This section includes:**

Supplementary Materials and Methods

Supplementary Note

Supplementary Figures S1 to S10

Supplementary Tables S1 to S3

Supplementary References

### Materials and Methods

#### 1 Strain construction

##### 1.1 Construction of Ptet driving *tetR* at *ycaD*, *intS*, and *galK* loci on chromosome

The *tetR* structure gene was amplified from pZS4int1 (*52*) by oligos tetR-Kpn-F and tetR-Bam-R (**Table S1b**). The PCR products were digested with *Kpn*I/*BamH*I and cloned into pKDT_Ptet (*53*) digested with the same enzymes, yielding pKDT_Ptet-*tetR*. Using the resultant plasmid as template, the DNA fragment (referred to as “km:rrnBT:Ptet-tetR”) containing the *km* gene, the *rrnB* terminator (rrnBT) and the Ptet promoter was individually amplified using three pairs of chimeric oligos: (1) Ptet.tetR-ycaD-F and Ptet.tetR-ycaD-R, (2) Ptet.tetR-intS-F and Ptet.tetR-intS-R, and (3) Ptet.tetR-galK-F and Ptet.tetR-galK-R (**Table S1b**). The PCR products were gel purified and individually integrated into the *ycaD*, *intS* and *galK* locations on the chromosome of K12 strain NCM3722 using the lambda-Red system (*54*). The Km resistant colonies were verified by PCR and subsequently by sequencing.

At the *ycaD* locus, the “km:rrnBT:Ptet-tetR” fragment is located in the *ycaC*/*ycaD* intergenic region, replacing the sequence from the -221^st^ nucleotide to the -114^th^ nucleotide relative to the translational start point of *ycdD*. The resultant strain is named NCM3722-1R. At the *intS* location, the “km:rrnBT:Ptet-tetR” fragment is substituted for the region from the -228^th^ nucleotide to the +1183^th^ nucleotide relative to the translational start point of *intS*. At the *galK* location, the “km:rrnBT:Ptet-tetR” fragment is substituted for the region from the -7^th^ nucleotide to the +1033^th^ nucleotide relative to the translational start point of *galK*. The three copies of Ptet-*tetR* were combined together by first flipping out the km resistance genes (*54*) and subsequently by P1 transduction, yielding strain NCM3722-3Rs (that is, HE697) (**Table S1a**).

##### 1.2 Construction of the *ackA/acs* deletion strain

The Δ*ackA* deletion allele in strain JW2293-1 (*E. coli* Genetic Stock Center), in which a *km* gene is substituted for *ackA*, was transferred by P1 transduction to NCM3722. The km resistance genes were then flipped out (*54*). The Δ*acs* deletion allele in strain JW4030-1 (*E. coli* Genetic Stock Center), in which a *km* gene is substituted for *acs*, was transferred by P1 transduction, yielding strain NQ1028 (**Table S1a**).

##### 1.3 Construction of the lacI*/lacZ/lacY* deletion strain

The *lacI*, *lacZ* and *lacY* genes were deleted using the lambda-Red method (*54*). The km resistance gene was amplified from pKD4 using chimeric oligos lacI-P1 and lacY-P2 (**Table S1b**). The PCR products were electroporated into NCM3722 cells expressing lambda-Red proteins encoded by pKD46 (*54*). The Km resistant colonies were confirmed by PCR and sequencing for the replacement of the region harboring *lacI*, *lacZ* and *lacY* genes by the *km* gene. This yielded strain NCM3722 Δ*lacIZY*. The *lacIZY* deletion was transferred by P1 transduction to NCM3722-1R, yielding strain HE827 (**Table S1a**). Plasmids pZA31Ptet-*lacY* (*1*) and pZA31Ptet-*gfp* (*55*) were individually transformed into HE827, yielding strain HE620 and strain HE829, respectively (**Table S1a**).

##### 1.4 Construction of Ptet driving *otsBA* on chromosome

Using plasmid pKDT:Ptet (*53*) as template, the DNA fragment (referred to as “km:rrnBT:Ptet”) containing the *km* gene, the *rrnB* terminator (rrnBT) and the Ptet promoter was amplified using the primer pair Ptet.ots-P1/Ptet.ots-P2 (**Table S1b**). The PCR products were integrated into the chromosome of K12 strain NCM3722 (*54*) to replace the *otsBA* promoter (from the -103^th^ nucleotide to the +1^st^ nucleotide relative to the translational start point of *otsB*). The chromosomal integration was confirmed first by colony PCR and subsequently by DNA sequencing. The region carrying “km:rrnBT:Ptet-otsBA” was transferred to NCM3722-1R by P1 transduction, yielding strain HE647 (**Table S1a**).

##### 1.5 Construction of *otsBA* overexpression plasmid

The *otsBA* operon was amplified from NCM3722 chromosomal DNA using oligos otsB-Kpn-F and otsA-Bam-R (**Table S1b**). Using fusion PCR, the *BamH*I restriction site GG**ATC**C (+383 to +388 downstream of the *otsA* translastional start point; ATC in bold face encoding the Isoleucine residue) on the *otsA* gene was removed by changing ATC to ATT. The modified *otsBA* operon with no *BamH*I site was digested with *Kpn*I/*BamH*I, gel purified and then ligated into the same sites of pZA31Ptet and pZE12Ptet (*52*), yielding pZA31Ptet-*otsBA* and pZE12Ptet-*otsBA*, respectively. pZA31Ptet-*otsBA* and pZE12Ptet-*otsBA* were transformed into strain HE697 (carrying three chromosomal copies of Ptet-*tetR*), yielding strain HE650(**Table S1a**).

##### 1.6 Construction of *araE* overexpression plasmid

The Δ*araC* deletion alleles in strain JW0063-1 (E. coli Genetic Stock Center), in which a *km* gene is substituted for *araC*, was transferred by P1 transduction to NCM3772-1R, yielding strain HE636. Because the parent strain contained was also deleted of *araBAD* (*56*), and the latter was adjacent to the *araC* locus, *ΔaraBAD* was also transferred to the recipient strain HE636.

The araE gene was PCR amplified from NCM3722 chromosomal DNA using oligos araE-Kpn-F and araE-Bam-R (**Table S1b**). The amplified products were digested with KpnI and BamHI, and then ligated into the same sites of pZA31Ptet (*52*), yielding pZA31Ptet-araE, in which the araE gene is driven by the tet promoter Ptet. pZA31Ptet-*araE* (*1*) and pZA31Ptet-*gfp* (*55*) were individually transformed into HE636, yielding strain HE639 and strain HE839, respectively (**Table S1a**).

##### 1.7 Construction of pHluorin expression plasmid

The pHluorin gene, encoding pH sensitive ratiometric GFP, was amplified from the plasmid pGFPR01 (*26*) using oligos pHgfp-EcoR-F and pHgfp-Bam-R (**Table S1b**). The amplified PCR products were digested with *EcoR*I and *BamH*I, purified and ligated into the same sites of pZE12Ptet, yielding pZE12Ptet-pHluorin. This recombinant plasmid was transformed into strain NCM3722-1R, yielding strain HE616 (**Table S1a**). The plasmid pZA31Ptet-*gfp* (*55*) was transformed into strain HE697, yielding the control strain HE828.

#### 2 Growth of Cells

##### 2.1 Growth media

The phosphate-based growth media contained 20 mM glucose 10 mM NaCl, 10 mM NH_4_Cl, 0.5 mM Na_2_SO_4_, a phosphate buffer and a 1000x micronutrient solution. The 1000x micronutrient solution contained 20 mM FeSO_4_, 500 mM MgCl_2_, 1 mM MnCl_2_·4H_2_O, 1 mM CoCl_2_·6H_2_O, 1 mM ZnSO_4_·7H_2_O, 1 mM H_24_Mo_7_N_6_O_24_·4H_2_O, 1 mM NiSO_4_·6H_2_O, 1 mM CuSO_4_·5H_2_O, 1 mM SeO_2_, 1 mM H_3_BO_4_, 1 mM CaCl_2_, and 1 mM MgCl_2_ dissolved in a 0.1 N HCl solution. The content of the phosphate buffer was changed to control the pH. At pH 6, the media was buffered with 20 mM K_2_HPO_4_ and 80 mM KH_2_PO_4_. The total osmolarity for the base medium including glucose at pH 6 is equal to 280 mOsm. For other pH, the proportion of K_2_HPO_4_ and KH_2_PO_4_ was used at different proportions with the total concentration summing to 100 mM. For internal potassium measurements, K_2_HPO_4_ and KH_2_PO_4_ was replaced with Na_2_HPO_4_ and NaH_2_PO_4_ and 1 mM KCl was added to provide some potassium.

The medium used for the anaerobic growth of B. thetaiotaomicron was the same as used for the anaerobic growth of E. coli but also included 2 mg cyanocobalamin, 2 mg hemin, and 0.6 cysteine per liter. To make the media anoxic, Hungate tubes (16 mm x 125 mm) filled with 7 mL medium were shaken at 270 rpm under a 7% CO_2_, 93% N_2_ atmosphere pressurized to 1.5 atm for 75 minutes. Cultures were transferred anoxically into Hungate tubes with disposable syringes.

The MOPS based growth media was the same as in Ref. (*57*). The base medium contains 40 mM MOPS and 4 mM tricine (adjusted to pH 7.4 with KOH), 0.1 M NaCl, 10 mM NH_4_Cl, 1.32 mM KH_2_PO_4_, 0.523 mM MgCl_2_, 0.276 mM Na_2_SO_4_, 0.1 mM FeSO_4_, and the trace micronutrients described in Ref (*58*). The MES buffered media was to the same as the MOPS media except the MOPS buffer and NaCl was replaced with 150 mM MES. In all media, carbon concentrations were added based on the number of carbon atoms in the molecule – 10 mM for C6 carbons, 20 mM for C3 carbons.

Ampicillin, chloramphenicol, and kanamycin were added at concentrations of 50 mg/mL, 10 mg/mL, and 50 mg/mL, respectively.

Stresses, limitations, and overdoses: Acetate stress was applied to cells by growing cells in minimal media with various concentrations of sodium acetate. Lactose overdose was applied to cells (HE620) by titrating lactose concentrations. For the same cultures, Chloro-tetracycline (cTc) was also added to the media at a concentration of 25 ng/mL to induce expression of LacY. Trehalose overdose was applied to cells (HE647 or HE650) grown in different concentrations of cTc (0-120 ng/mL). CCCP stress was applied for cells (HE616) by titrating CCCP between 0-40 µM. Carbon limitation was implemented by titrating 3-methyl-benzylalcohol (3MBA) concentration in strains NQ1243 and NQ1390. 3MBA concentrations ranged from 0 to 600 µM.

##### 2.2 Measurement of pH

Media pH was measured with an Orion Start A221 pH meter by Thermo Scientific with an Orion 9110DJWP pH probe by Thermo Scientific. Measurements were made according to the manufacturer’s instructions.

##### 2.3 Culture tubes

Exponential cell growth was performed in a 37°C water bath shaker at 240 rpm Cultures were grown in the following three steps: seed culture, pre-culture, and experimental culture. Cells were first grown as seed cultures in LB broth for several hours, then as pre-cultures overnight in an identical medium to the experimental culture. Experimental cultures were started by diluting the pre-cultures to an optical density (OD) at wavelength 600 nm (OD_600_) of ∼0.01–0.02. Growth rates were calculated from at least seven OD_600_ points within a range of OD_600_ of ∼0.04–0.4.

##### 2.4 Plate reader

Seed culture and pre-culture were performed in water bath shakers as described for the growth of cells in culture tubes. Experimental culture was done in a Tecan Spark microplate reader with 96-well microplates (Greiner bio-one) with 200 µL of media. For inoculation, cells were diluted at least 1,000x into the plate media. The incubation temperature was 37°C. The plate was shaken at 280 rpm. Optical density was measured at a wavelength of 420 nm (OD_420_). To calculate growth rate, the background OD from the opacity of the plate and media, was subtracted from raw OD measurement. Growth rates were calculated from OD_420_ from 0.02-0.2. Fluorescence was also measured for internal pH (see internal pH – fluorescence).

##### 2.5 Anaerobic cultures

Anaerobic growth was performed similarly to aerobic growth with a few exceptions. All transfers were performed with disposable syringes to avoid oxygen contamination. For *E. coli*, aerobic seed cultures were diluted into Hungate tubes for preculture. After overnight growth, the precultures were diluted into fresh Hungate tubes for experimental culture. For B. theta, the seed cultures were inoculated into Hungate tubes containing 7 mL Wilkens-Chalgren broth from colonies selected from Wilkens-Chalgren agar plates. After overnight growth, these cultures were diluted into preculture tubes. And then diluted once more for the experimental cultures. To avoid atmospheric exposure from removing samples, OD measurements were performed with a Thermo Genesys 20 modified to hold Hungate tubes in place of cuvettes. The culture temperature was kept stable during OD measurements by removing and replacing the Hungate tubes from the water bath shaker within 30 seconds. The OD_600_ measured through the Hungate tubes was equivalent to the OD_600_ measured through a cuvette for the range of 0.04-0.5.

Anerobic growth in plate reader was performed with a Tecan Spark microplate reader enclosed in a custom vinyl anaerobic chamber. Chamber was kept anaerobic with palladium catalysts and an input gas of 5% H_2_, 10% CO_2_, and 85% N_2_. Oxygen levels were monitored with the Tecan Spark O_2_ and CO_2_ module to ensure the chamber stayed anaerobic during growth.

#### 3 Assays

##### 3.1 RNA measurement

Total RNA quantification method used was described in Ref (*35*). RNA nucleotide amount (nmol/OD/mL) was calculated from the normalized RNA mass (µg/OD/mL) assuming that the average molecular weight of an RNA nucleotide was 339.5 g/mol, equivalent to equal parts ACGU.

##### 3.2 Internal acetate, lactose, and trehalose measurements

Exponentially growing cells were harvested at an OD of 0.4-0.5. 600 µL of culture was collected and added to a 0.22 µm nylon filter centrifuge tube (Corning Costar Spin-X Centrifuge Tubes) and centrifuged at 20,000g for 30 seconds. Cells were quickly removed from the filter with 600 µL of deionized water. The cell-free filtrate was put aside for later. The extracted cells were then added to a 1.5 mL Eppendorf centrifuge tube containing 40 µL chloroform. The tube was vortexed for 10 seconds. Samples were stored at −20°C. For the cell-free filtrate, the same procedure was applied, starting by repeating the filter step for the filtrate.

For HPLC analysis, samples were thawed and then centrifuged for 60 seconds. 80 µL of the aqueous section was placed into HPLC analysis tubes for analysis with HPLC (Ref (*4*)).

The internal acetate or lactose amount, [*Ac*]*_internal_*, was calculated as 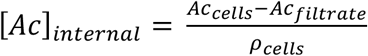 where Ac_cells_ is the acetate amount in the filtered cells, Ac_filtrate_ is the acetate measured in the re-filtered filtrate, and ρ_cells_ is the amount of cells analyzed (in ODmL). When concentration was reported in mM, the formula was 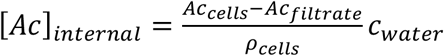 where c_water_ was assumed to be 1 µL H_2_O/OD/mL.

Trehalose pools were measured as follows. Exponentially growing cells were harvested at an OD of 0.4-0.5. A 1 mL of culture was collected on a membrane filter (Durapore membrane filters; 0.45 um; HVLP02500) which was set on a vacuum manifold and prewashed by water followed by warmed culture medium. The cells were rinsed by 2 mL warmed culture medium and transferred to 2 mL of extraction solution (40:40:20 acetonitrile:methanol:water) containing 100 nmol glycerol and 50 nmol arabinose as standards placed on a dry ice plate. Cells on the filter were suspended well in the extraction buffer by pipetting and extraction was let proceed at −20°C for 40-60 minutes. After the extracts were collected to microtubes, the filter was rinsed by 0.9 mL extraction solution and the extraction was let proceed for another 15-20 minutes. This second extracts were combined with the first extracts and stored at −80°C. On the day of HPLC measurements (Ref (*4*)), the extracts were dried by speed-vac and dissolved in water.

##### 3.3 Excreted glucose and acetate

Four samples of 200 µL were pipetted from culture tubes at regularly spaced ODs during exponential growth. For anoxically grown cultures, sample removal was done with tuberculin syringes inserted into the rubber stopper. Samples were transferred to 0.22 µm nylon filter centrifuge tubes (Corning Costar Spin-X Centrifuge Tubes) and quickly filtered by centrifugation. Samples were then stored at −20°C until HPLC analysis, which was performed according to Ref (*4*).

##### 3.4 Internal glutamate measurement

Measurement of amino acids were performed as in Refs. (*59–61*). Briefly, 150 µL of exponentially growing cells were added to 600 µL of ice-cold methanol with alpha amino-butyric acid as an internal standard. The samples were vortexed and then kept at 4°C for storage. The samples were then dried in a rotary vacuum desiccator and resuspended in 150 µL of H_2_O for HPLC analysis. Glutamate levels were normalized by the glutamate level for unstressed cells.

##### 3.5 Potassium

Cells for potassium measurements were grown in low potassium media where the potassium salts were replaced with sodium. 1 mM of KH_2_PO_4_ was used as the source of potassium. Exponentially growing cells were harvested at OD_600_=0.4-0.5. 2 mL of culture was collected on a membrane filter (Durapore membrane filters; 0.45 µm; HVLP02500) set on a vacuum manifold. The cells were rinsed by 2.5 mL warmed potassium-free medium and transferred to 1.8 mL of 1 N HNO_3_.

Cells on the filter were suspended well in the extraction buffer by pipetting and extraction was let proceed at ambient temperature for 30 minutes. After the extracts were collected to microtubes, the filter was rinsed by 1 mL of 1 N HNO_3_ and the extraction was let proceed for another 15 minutes. This second extracts were combined with the first extracts and stored at −80°C. Potassium was measured in these samples with ICP-MS at the Environmental and Complex Analysis Laboratory (ECAL) at UCSD.

##### 3.6 Protein measurement

Total protein quantification method used was described in Ref (*35*). Bovine serum albumin as used as the protein standard.

#### 4 Metabolomics

##### 4.1 Sample collection

2.5 OD_600_·mL of exponentially growing cell culture were quickly vacuum filtered on 0.45 µm nylon filters. The cells on the filter were quickly washed 2 times with 2 mL of warm cell-free media. After washing, the filter was immediately plunged into 1.3 mL of a 40:40:20 methanol:acetonitrile:water (MAW) mixture kept on dry ice for 15 minutes. The MAW mixture without the filter was then put into a 1.5 mL Eppendorf tube and centrifuged for 60 seconds. The supernatant was put into a new tube and stored at −80°C.

To perform relative quantification with ^13^C-labeled metabolites, cells were grown in media supplemented with ^13^C carbon sources were collected for use as a reference. A mixed reference was used to avoid bias from any of the individual growth medium. Three conditions were collected: NCM3722 cells without an applied stress, NCM3722 cells grown with 30 mM ^13^C-labeled sodium acetate, and HE650 cells grown with 50 ng/mL cTc (high level of trehalose overdose, see **Fig. S9**). All conditions were grown in phosphate-buffered minimal media with ^13^C-labeled glucose as the carbon source. Sample collection was the same as described above. Samples were combined in equal amounts to form the mixed ^13^C-labeled reference.

Extracted metabolites were analyzed by LC-MS. For hydrophilic interaction chromatography, used a Vanquish UHPLC system (Thermo Fisher, San Jose, CA) and an XBridge BEH Amide column (2.1 mm x 150 mm, 2.5 mm particle size, 130 A ° pore size; Waters, Milford, MA) with a 25 min solvent gradient at flow rate of 150 µL/min. Solvent A is 95:5 water:acetonitrile with 20 mM ammonium hydroxide and 20 mM ammonium acetate, pH 9.4. Solvent B is acetonitrile. The LC gradient was 0 min, 85% B; 2 min, 85% B; 3 min, 80% B; 5 min, 80% B; 6 min, 75% B; 7 min, 75% B; 8 min, 70% B; 9 min, 70% B; 10 min, 50% B; 12 min, 50% B; 13 min, 25% B; 16 min, 25% B; 18 min, 0% B; 23 min, 0% B; 24 min, 85% B; 30 min, 85% B. LC was coupled to was a quadrupole-orbitrap mass spectrometer (Q Exactive, Thermo Fisher Scientific, San Jose, CA) via electrospray ionization. The mass spectrometer operates in negative and positive ion switching mode and scans from m/z 70 to 1000 at 1 Hz and 70,000 resolution. Autosampler temperature was 5°C, and injection volume was 10 µL. Data were analyzed using the El-Maven software. ^13^C natural isotope abundance was corrected using house code (*62*).

##### 4.2 Relative quantitation

Relative quantities were calculated as 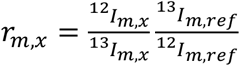 where r_m,x_ is the relative quantity of metabolite m in condition x, and ^n^I_m,x_ is the ion count for metabolite m with atomic mass of carbon n (12 or 13) in either condition x and ref being the reference condition. r_m,x_ is the relative value of metabolite m in condition x, and r_m,ref_ is the relative value of metabolite m in the reference condition. The reference condition consisted of uninhibited samples from each limitation or stress.

##### 4.3 Absolute quantitation

To compute absolute quantities of metabolites, we use the formula [*M*]*_x_*_,*m*_ = [*M*]*_p_*_,*m*_*r*_*m,x*_ where [M]_x,m_ is the concentration of metabolite m in condition x, [M]_p,m_ is the concentration of metabolite m as measured in Ref (*32*) for their glucose condition.

When calculating the summed metabolite concentration including free potassium, the concentrations of glutamate and aspartate were multiplied by 2 to account for potassium associated with these negatively charged molecules.

#### 5 Internal pH

##### 5.1 Fluorescence

Fluorescence measurements were taken for exponentially growing HE616 cells harboring constitutive expression of pHluorin in the OD range of 0.01 and 0.2 using a Tecan Spark microplate reader. Fluorescent signals were collected for two excitation wavelengths: 390nm and 470nm. The emission wavelength for both signals was 500 nm. For each growth condition, a linear fit was made from the two signals to calculate the fluorescence ratio. This fluorescence ratio was then used to calculate the internal pH from the standard curve. In cells grown with CCCP, the fluorescence signal was adjusted to account for light absorbance from CCCP.

The following procedure was used to make the standard curve. HE616 cells grown in phosphate buffered media were resuspended in carbon-free phosphate-buffered media with 10 µM CCCP for a variety of pH. Using a Tecan Spark microplate reader, fluorescence with excitations at 390 nm and 470 nm was measured for cells at different dilutions. The fluorescence emission wavelength for both signals was 500 nm. For each pH, a linear fit was made from the two signals. The slope from this fit was used as the fluorescence ratio.

##### 5.2 Radiolabel

Adapted from Refs. (*27, 28*). Cells were grown in 150 mM MES medium, pH 6.3, to OD_600_ = 0.5 and a 3.2 mL of culture was transferred to a tube containing 66 μL of ^14^C-sucrose (0.25 mM, 50 μCi/mL from Perkin Elmer Inc.) and 33 μL of either [^3^H]-benzoic acid (0.5 mM, 0.5 mCi/ml from ARC) or ^3^H-H_2_O (1 mCi/mL from Perkin Elmer Inc.) and incubated at 37°C for 2-3 minutes. A 1 mL suspension was transferred to a microtubes containing 200 μL of 1-bromododecane and centrifuged at 21130 x g for 30 seconds. The supernatants were carefully removed, and the cells were suspended in 100 μL water. The supernatant and the cell suspension were transferred to 15 ml of scintillation cocktail (Liquiscint from National Diagnostics) and analyzed by scintillation counter. ^14^C counts into ^3^H channel were corrected. Impurities in ^14^C-sucrose and ^3^H-benzoic were removed by suspending exponentially growing NCM3722 cells followed by centrifugation and taking the supernatants. Cytoplasmic pH was calculated from the ratio of the concentrations of benzoic acid between the cytoplasm and the supernatant.

##### 5.3 Cytoplasmic water

Adapted from Ref (*63*). For cytoplasmic water measurements under carbon limitation and trehalose overdose, cells were grown in MOPS-buffered medium. The culture was processed essentially as described above with ^14^C-sucrose and ^3^H-H_2_O except that oil centrifugation was done for 3 min.

#### 6 Proteomics

Proteomics samples were taken for acetate stressed cells and lactose overdose cells. Samples were prepared as in Ref (*64*) with the following exceptions. For the ^15^N reference, unstressed cells growing in glucose, phosphate-based media (**Methods 2.1**) where the NH_4_Cl was replaced with 10 mM ^15^NH_4_Cl.

Proteomics for glutamate synthesis limitation, sugar uptake limitation, and protein synthesis limitation were adapted from (*29*). Because only the relative data was provided in that publication, protein abundances were converted to mass fraction by matching the mass fractions for each gene from the condition with no additional acetate for the acetate-stress series (**Table S2a**).

### Supplementary Note

As described in the main text and **Fig. 4b**, the accumulation of potassium acetate in the cell is controlled by the internal pH. Potassium acetate accumulation itself exerts an inhibitory effect on cell growth as a useless metabolite. This effect can be reduced by lowering the internal pH. But the pH reduction independently reduces growth rate as well (**Fig. S1f**). Here we develop a simple model to study the consequence of this tradeoff.

The reduction of growth from useless metabolites is described by Eq. (3) and **Fig 3f**. For a useless metabolite at concentration *m*, we write the growth rate as *λ* = *λ*_0_ ⋅ *β*(*m*), where *λ*_0_ is the unstressed growth rate and

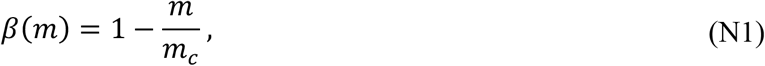

with *m_c_* being the inhibitory concentration where growth vanishes. The value of *m*_c_ is not known for potassium acetate as a useless metabolite, but we will estimate it later. When the useless metabolite is potassium acetate, the intracellular concentration is dependent on the internal pH through the Henderson-Hasselbalch equation (Eq. 1),

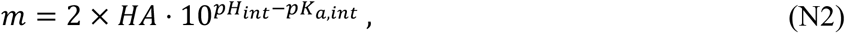

where *HA* is the concentration of acetic acid which has equilibrated across the cell membrane, *pH_int_* is the internal pH value of the cell, and *pK_a,int_* ≈ 4.97 is the effective pKa value in the cell according to the data in **Fig. S3d**. A factor of 2 is included to account for the potassium associated with acetate.

When cells experience a reduction in internal pH (without useless metabolite accumulation), they also suffer a growth reduction. We describe the data in **Fig. S1f** with a quadratic form, *λ* = *λ*_0_ ⋅ *γ*(*pH*), where

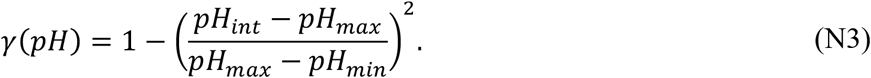

Here *pH_max_* is the internal pH value of unstressed cells, and *pH_min_* is the internal pH at which growth stops. **Fig. S10a** shows that the data is well described by the form in Eq. (N3), with the values *pH_max_* = 7.78, *pH_min_* = 6.42, and 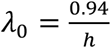.

To model the combined effect of potassium acetate and internal pH, we have to make an assumption on the function form of the growth rate *λ*: If *m* = 0 so that *β* = 1, then *λ* = *λ*_0_ ⋅*γ*(*pH_int_*) and if *pH_int_* = *pH_max_* such that *γ* = 1, then = *λ*_0_ ⋅ *β*(*m*). Here we assume the joint effect to be described by a simple product form, which is the simplest form satisfying the above constraint,

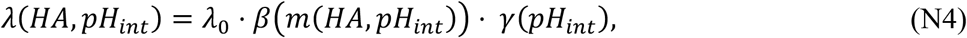

where *m*(*HA*, *pH_int_*) is given by Eq. (N2).

The only unspecified variable in Eq. (N4) is the internal pH, *pH_int_*, which is under the control of the cell. For a fixed stress level imposed by an external acetic acid concentration *HA*, the growth rate given by Eq. (N4) has a unique maximum in the range *pH_min_* < *pH_int_* < *pH_max_* (see **Fig. 4c**). In our model, we assume that the cell sets the internal pH at a level 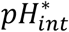 which maximizes the growth rate. The value of 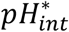 is obtained by setting 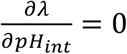, where*λ*_0_ comes from Eq. (N4):

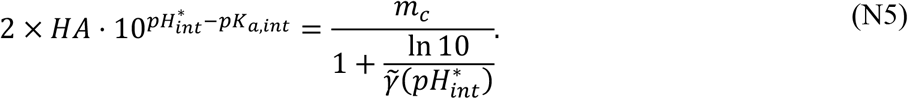

where

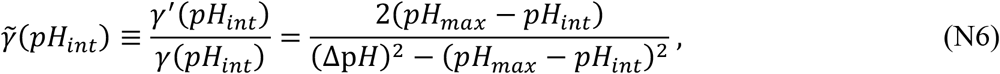

and Δp*H* ≡ *pH_max_* – *pH_min_* ≈ 1.36.

As mentioned already, we do not know the parameter *m_c_* for potassium acetate as a useless metabolite. We can fix this parameter by fitting the relation between internal pH and the internal potassium acetate concentration found by substituting Eq. (N2) into Eq. (N5). Denoting the internal acetate concentration corresponding to the value of *pH*^∗^ by *m*^∗^, we obtain:

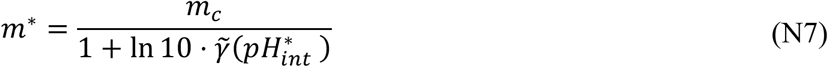

This relationship is plotted in **Fig. S10b** (solid line). The data is well described for *m_c_* ≈ 620 *mM*.

Notably, the relation between *m*^∗^ and 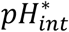 is approximately linear over its range. So, we explored a linear solution as well. To do so, let us simplify the notation first: Let 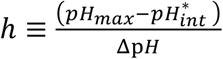 and let *x* = *γ̃* ⋅ Δp*H*. Then Eq. (N7) is a quadratic equation for ℎ(*x*), i.e., *x* ⋅ (1 − ℎ^2^) = 2ℎ, with the solution

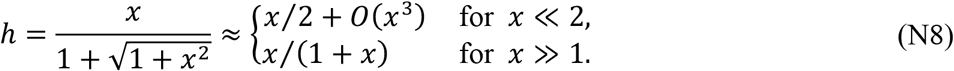

Further using 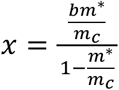 from Eq. (N7) where *b* ≡ Δp*H* ln 10 ≈ 3.1, we obtain

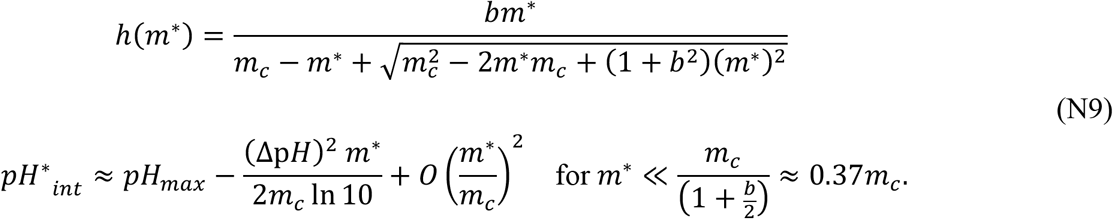

The linear (small-x) solution in Eq. (N9) is plotted in **Fig. S10b** as a dashed line. We see that the exact solution is well approximated by the small-x solution (dashed line) over the regime where data is available. The linear form of the pH-acetate relation shows that for much of the data range, the inhibitory concentration *m_c_* is shifted to a smaller value, 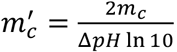, which is the intersection of the dashed and dotted line in **Fig. S10b**, where *pH*^∗^ → *pH_min_* (or ℎ → 1) and growth rate vanishes. From the best-fit parameter *m_c_* ≈ 620 *mM*, we obtain 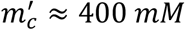. The difference between the two parameters reflects the additional cost due to pH reduction.

Using Eq. (N9), we can express the growth rate 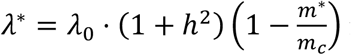 in term of the internal potassium acetate alone. **Fig. S10c** shows that the approximate linear decrease of growth rate against the internal potassium acetate concentration, shown also in main text **Fig. 3f**, recovered here as the small-x expansion. The dotted line is obtained using the approximate linear relation between *m*^∗^ and ℎ^∗^, i.e., for

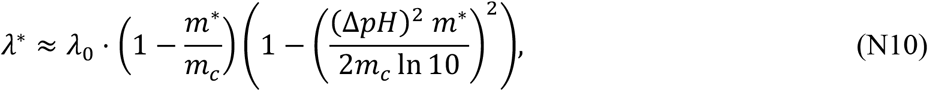

which has *λ*^∗^ → 0 as 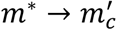, the apparent inhibition concentration.

The exact relation between growth rate and internal pH, plotted as the solid line in **Fig. S10d**, is obtained from substituting the metabolite concentration from Eq. (N7) into Eq. (N4) as

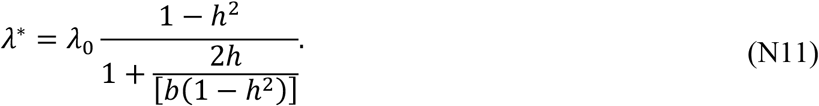

Finally, the dependence of the growth rate on the acetic acid concentration, shown as the solid line in **Fig. 4b**, is obtained as an implicit function defined by *λ*^∗^(ℎ) in Eq. (N11) and *HA*(ℎ) from Eq. (N5):

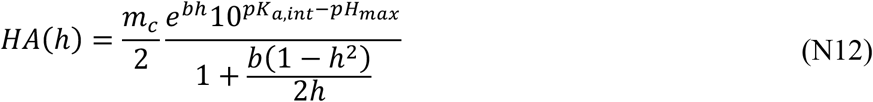

We can ask what parameters the cell can change to increase tolerance to acetate. To do this, we solve the half-inhibitory acetic-acid concentration, *HA*_50_, which is the acetic acid concentration where the growth rate is halved. To find *HA*_50_, first we solve the value ℎ_50_(*b*) where *λ*^∗^ = 0.5*λ*_0_ using Eq. (N11), and then insert ℎ_50_(*b*) into Eq. (N12). For 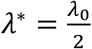, we find 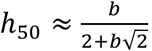. Substituting this dependence of ℎ_50_(*b*) into Eq. (N12), we obtain

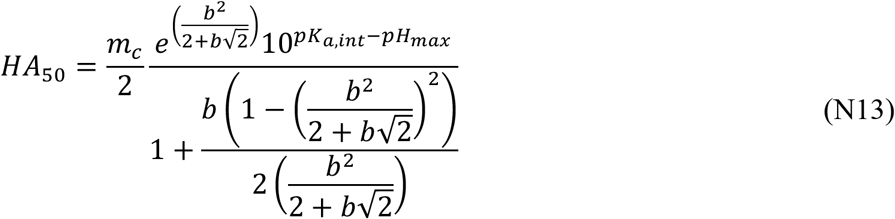

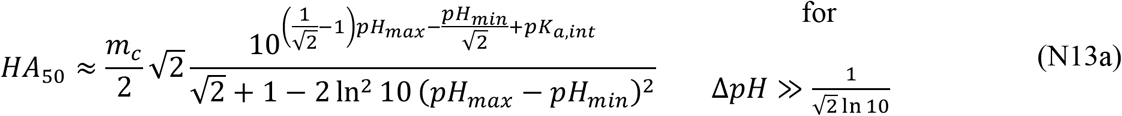

Eq. (N13) and (N13a) show that up to a substance-specific effect represented by *m_c_*, the half-inhibitory concentration *HA*_50_ is dependent on both the value of the stress-free pH (*pH_max_*) and the minimal pH (*pH_min_*), with 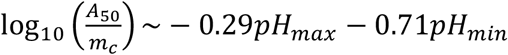. Thus *HA*_50_ increases (i.e., the cell becomes more tolerant to acetate) with either decrease in the normal (max) pH or decrease in the minimal pH, but with the latter being about twice as potent as the former. The full dependence of 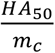 on *pH_max_* – *pH_min_* and on *pH_max_*, based on the exact solution, is shown in **Fig. N1a**, and the dependence on *pH_min_* and *pH_max_* is shown in **Fig. N1b**. Contour plots are shown in **Fig. S10e,f**.

**Figure N1.**
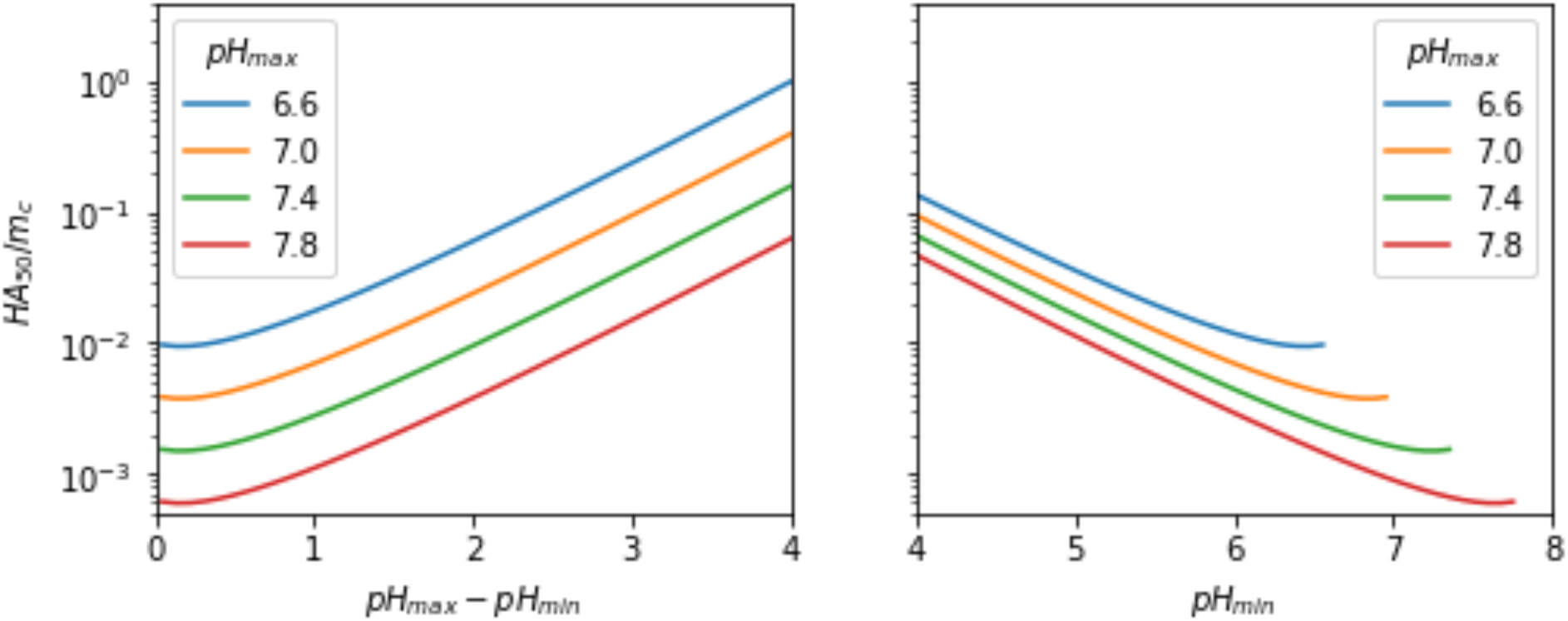
The dependence of the half-inhibitory acetic acid concentration *HA*_50_ on model parameters. **a)** *HA*_50_ expressed in term of the substrate-specific characteristic concentration *m_c_* given by Eq. (N13), for various values of stress-free pH (p*H_max_*) and the viable pH range *pH_max_* – *pH_min_*. **b)** Same quantity as that plotted in panel b, but for various values of the stress-free pH (*pH_max_*) and the minimal pH (*pH_min_*).

### Supplementary Figures

**Figure S1.**
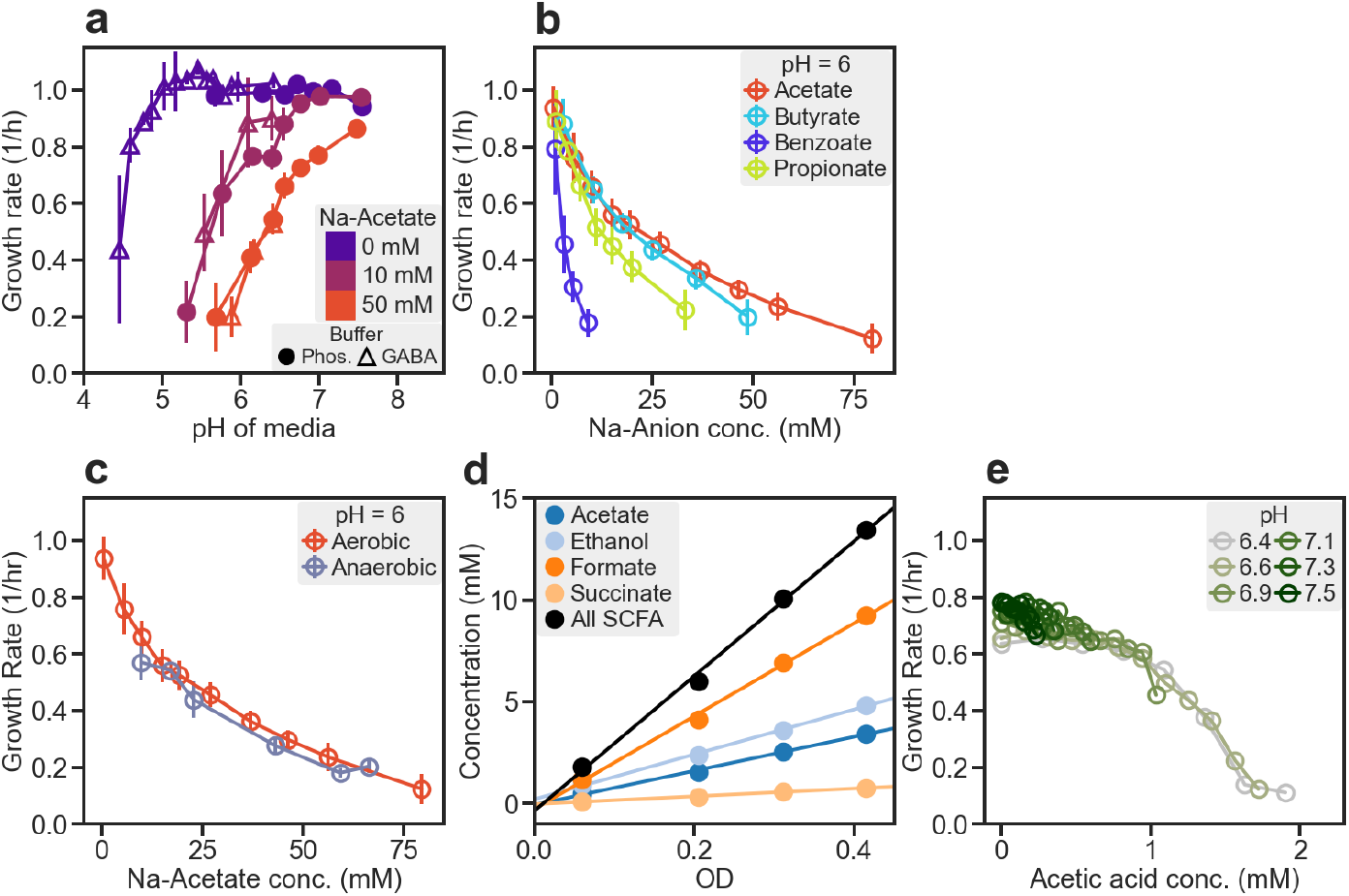
Response of *E. coli* to acetate and other SCFAs. Unless otherwise indicated, *E. coli* K-12 NCM3722 cells were grown exponentially in glucose minimal media, at various pH fixed by phosphate buffer. Cultures were grown in either test tubes or Tecan microplate reader; see **Methods 2**. **a)** Phosphate- or GABA-based buffers (circles or triangles, respectively) were adjusted to obtain a range of medium pH; see **Methods 2.1**. Blue, purple, red symbols represent cultures supplemented with different concentrations of sodium acetate (NaAc) Added to the medium. **b)** Growth in medium fixed to pH 6 and supplemented by various weak organic acids. Each acid was added as the sodium salt at the concentration indicated on the x-axis. **c)** Cultures were grown anaerobically in Hungate tubes with an atmosphere of 7% CO_2_ and 93% N_2_, with pH fixed to 6; see **Methods 2.5**. Various amounts of sodium acetate were added to the medium as indicated by the x-axis. Because large amounts of SCFA (totaling ∼15 mM, see panel d) was excreted over the course of growth, growth rate was not well-defined for medium with <15 mM sodium acetate added. Results of aerobic growth (red symbols) are shown for comparison. **d)** Measurements of various SCFA concentrations excreted by cells growing anaerobically in glucose minimal medium. Cultures were grown in Hungate tubes with an atmosphere of 7% CO_2_ and 93% N_2_, with pH set to 7; see **Methods 2.5** and **3.3**. **e)** Effect of pH and acetate on *B. thetaiotaomicron* (ATCC 29148), grown in glucose minimal medium with various amounts of sodium acetate, in phosphate buffered medium set to indicated pH; see **Methods 2.1** and **Table S1a**. (The difference in pH before and after growth was less than 0.15. The total amount of SCFA excreted was ∼5 mM, much smaller than anaerobically grown *E. coli* (panel d) due to the lack of formic acid excretion.) Anerobic growth was performed in microplate reader enclosed in a custom vinyl anaerobic chamber. Data is plotted against the acetic acid concentration, calculated by Eq. (1). Data in panels a, b, and c are binned for similar x-axis values, with the bins containing data from n ≥ 3 biological replicates. Error bars are calculated from the standard deviation.

**Figure S2.**
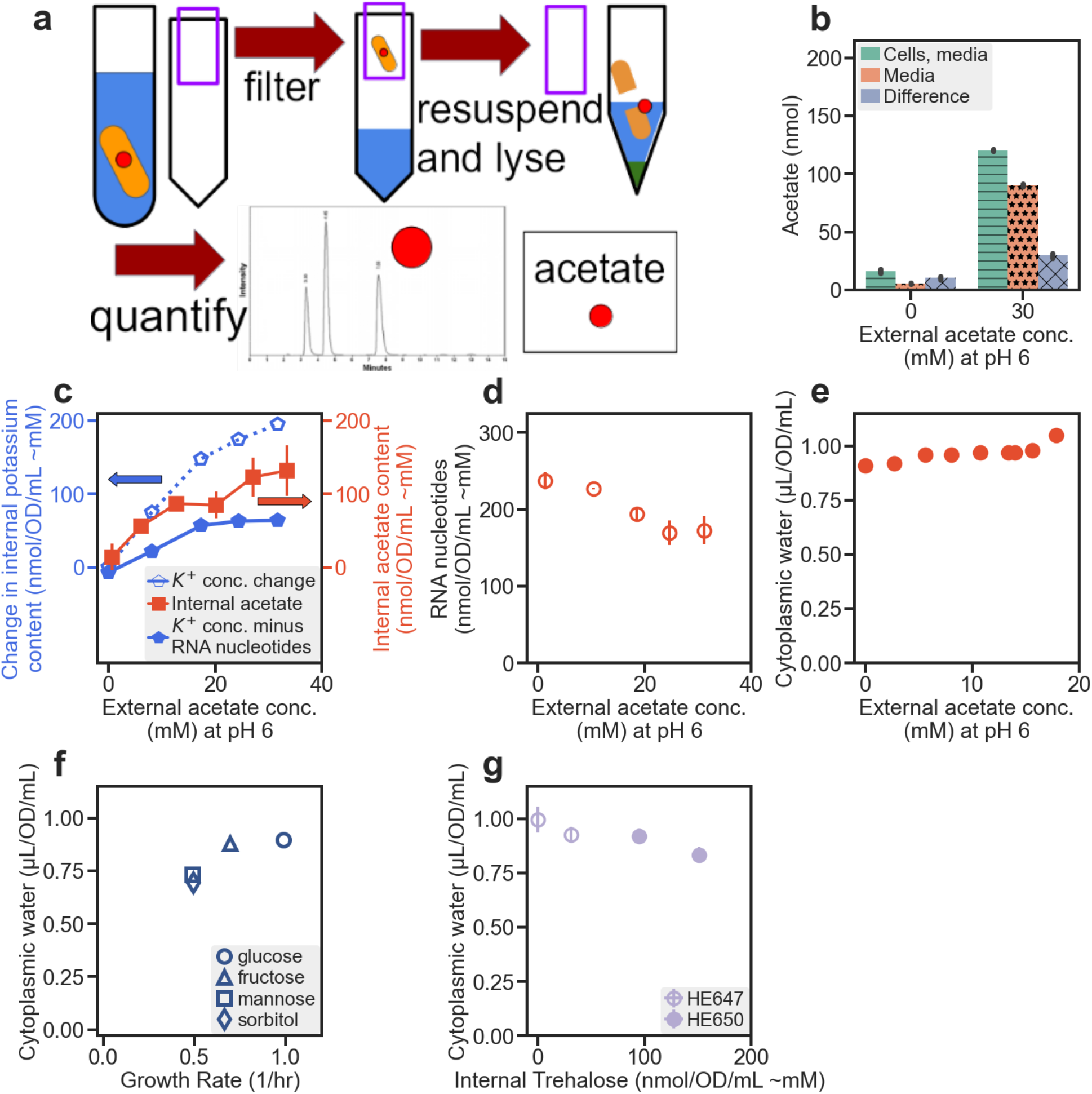
Internal acetate and water content measurements. **a)** Schematic of internal acetate measurement. Cells grown in test tubes were quickly centrifuged and filtered (< 30 sec) from steady state growing cultures to separate cells from the supernatant. Supernatant-free cells were then resuspended in distilled water and lysed with chloroform. The aqueous section was collected for analysis with HPLC. To correct for the acetate left on the filter from the residual media, the same procedure was performed again with the cell-free filtrate. **b)** Internal acetate: subtraction of media component. Cells and cell-free filtrate were collected as described in panel a. Reported in this panel are examples for cells grown with 0 and 30 mM sodium acetate added to the media. Acetate amount measured from the filtered cells are shown as green bars with horizontal lines, and those from the filtered media are shown as orange bars with stars. The internal acetate amount is obtained from the difference of the two and shown as purple bars with crossed lines. The acetate abundance (in unit nmol/OD/ml) was obtained by normalizing by the amount of cells (in OD·mL) added to the filter. **c)** To check for charge neutrality, the cellular potassium content was measured using ICP-MS (See **Methods 3.5**); change in the total potassium content is plotted against the external acetate concentration as solid blue symbols. Although the increase in total potassium filled blue symbols) was below the increase in internal acetate (red squares), additional potassium would be released as free ions due to the decrease in RNA content (panel d), because potassium is one of the main positive counter ions neutralizing the negatively charge phosphate backbone of RNA. The open blue symbols show the total increase in the free potassium ion pool, assuming that the reduction of each nucleotide residue in RNA leads to the addition of one free potassium ion. Our data suggests neutralization of acetate involves 0.5 potassium per nucleotide of the ribosome, consistent with the measured ratio of phosphate to magnesium, the other major counter ion stabilizing the ribosome. **d)** RNA content (**Methods 3.1**) of acetate stressed cells grown in phosphate buffered glucose minimal media at pH 6. **e)** Cytoplasmic water measurement in acetate stressed cells. Acetate stressed NCM3722 cells were grown in MES-buffered media at pH 6.3. Cytoplasmic water content was measured with radiolabeled ^3^H-water and ^14^C-sucrose (*63*), after they were added to steady state cultures and allowed to incubate for a brief period. The radioactivity of the two isotopes was measured from cells collected by centrifugation. Cytoplasmic water volume was calculated from the difference between estimated sucrose (extracellular + periplasmic) and water (extracellular + periplasmic + cytoplasmic). The same method was used to quantify the water content in panels f and g. **f)** Cytoplasmic water content for NCM3722 cells grown in MOPS minimal medium (pH 7.4) with various carbon sources. **g)** Cytoplasmic water content for cells accumulating internal trehalose; see **Fig. S9a**. Data in panels b, c (Internal acetate only), d, and g are grouped according to the same applied stress level and contain n ≥ 3 biological replicates. Error bars are calculated from the standard deviation.

**Figure S3.**
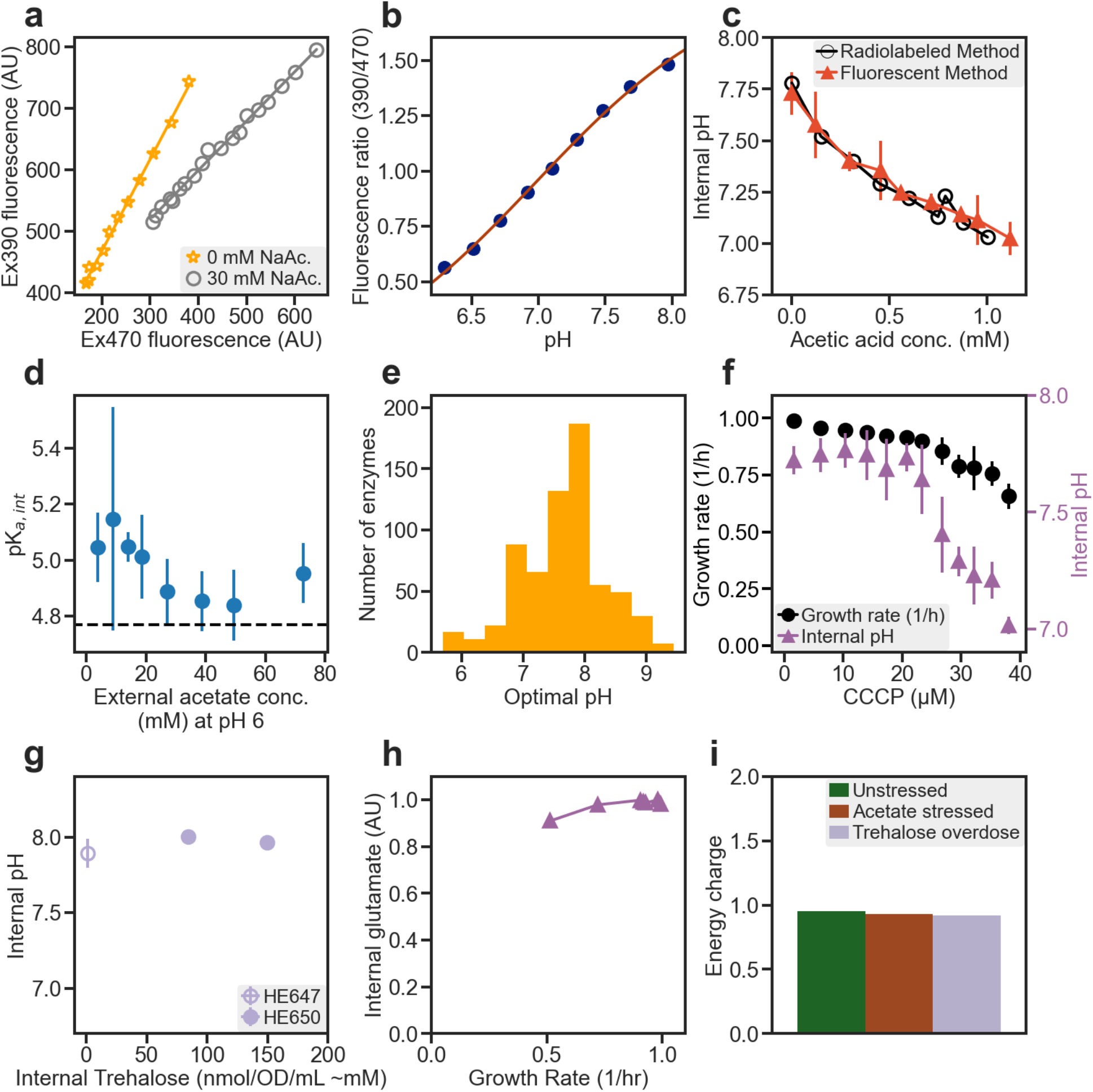
Measurement of internal pH. **a)** Fluorescence measurements were taken for exponentially growing HE616 cells harboring constitutive expression of ratiometric pHluorin (*26*) in the OD range of 0.01 and 0.2 using a Tecan Spark microplate reader. The fluorescent signals were collected for two excitation wavelengths: 390nm (y axis) and 470nm (x axis). Shown in the plot are data for two growth conditions, indicated by orange stars and grey circles (0 and 30mM sodium acetate, respectively, at pH 6 in phosphate buffered glucose media.) For each growth condition, a linear fit was made from the two signals to calculate the fluorescence ratio. This fluorescence ratio was then used to calculate the internal pH from the standard curve. **b)** Ratiometric pHluorin standard curve. HE616 cells grown as described in panel a were resuspended in carbon-free phosphate-buffered media fixed to a variety of pH. 10 µM CCCP was added to equilibrate the internal and external pH. For each pH, fluorescence ratio (390nm/470nm) was obtained as described in panel a for cells diluted to different densities. (The fluorescence signals settle down 15 min after CCCP addition, and the value of ratio reported here was taken an hour after CCCP addition.) **c)** Comparison of two internal pH measurement methods: by radiolabeled benzoic acid (black open circles – NCM3722) and by fluorescence with pHluorin (red triangles - HE616). Acetate stressed cells were grown in either MES-buffered media (radiolabeled method) at pH 6.2 or phosphate-buffered media (fluorescence method) at pH 6. Acetic acid concentration was calculated from the Henderson-Hasselbalch equation assuming a pKa of 4.77 in the medium. Internal pH for the radiolabeled method was found by measuring the fraction of radiolabeled ^14^C-benzoic acid taken up by exponentially growing cells (*27, 28*). Data for the fluorescence method is taken from **Fig. 1e** for comparison. **d)** For cells grown with varying concentrations of sodium acetate as shown in **Fig. 1d**, we define the intercellular pK_a_ by the Henderson-Hasselbalch equation, with 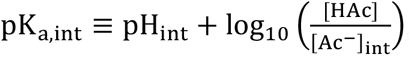, using the measured internal acetate concentration (**Fig. 1d**) and the measured internal pH (**Fig. 1e**). To bridge between data collected from separate experiments, internal pH and internal acetate concentrations were binned according to external acetate concentrations. Vertical lines represent the propagated standard deviation for the binned values. The average shift in the calculated pK_a,int_ from the accepted value of 4.77 (dashed horizontal line) was noted as a small difference in the predicted and observed acetate concentration in Ref. (*65*). **e)** Distribution of optimal pH for 664 enzymes in BRENDA database (*66*). **f)** Growth rate and internal pH of HE616 cells grown in glucose medium and no acetate, with various supplement of CCCP. pH of the medium was fixed to 6.0 by phosphate buffer. Same results were obtained for medium pH fixed to 7.0. **g)** Growth rate vs internal pH for HE647 and HE650 cells growing in phosphate buffered glucose medium with various levels of trehalose accumulation; see **Fig. S9a**. **h)** Internal glutamate concentration measured in CCCP limited cells. Cells grown in phosphate buffered media at pH 6 with varied concentrations of CCCP to reduce growth rate. Samples were collected with the no harvest protocol and glutamate content was measured with HPLC. **i)** ATP charge ratio was calculated as ([*ATP*] + .5 ⋅ [*ADP*])/([*ATP*] + [*ADP*] + [*AMP*]) from metabolomics measurements (**Table S3a**). The standard deviation of the energy charge in all conditions was less than 1%. Data in panels c (fluorescent method only), d, f, and g are grouped according to the same applied stress level and contain n ≥ 3 biological replicates. Error bars are calculated from the standard deviation.

**Figure S4.**
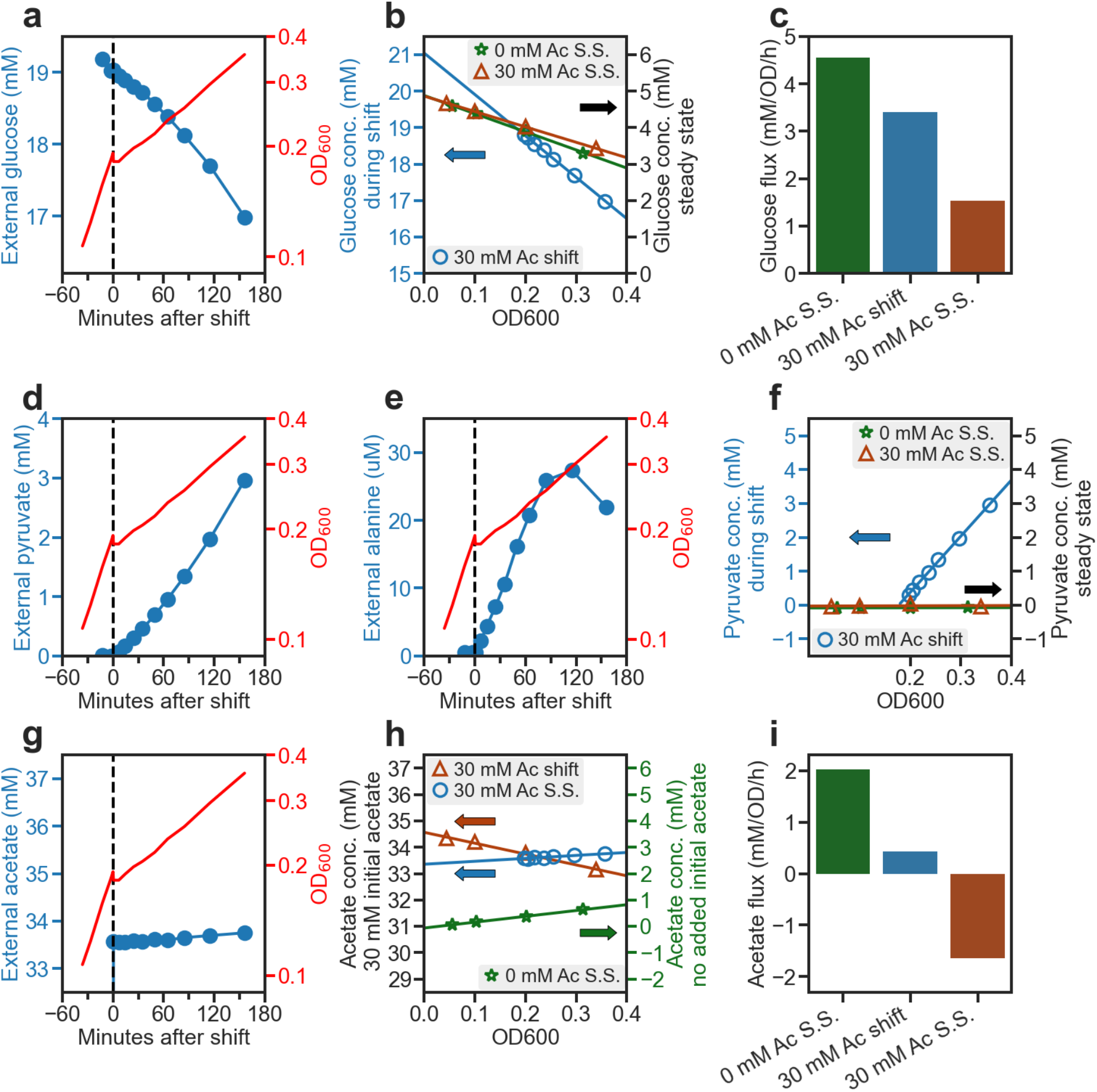
Extended disruption of carbon utilization following acetate addition. Unlike the quick adaptation in growth rate and glutamate/glutamine pools (**Fig. 2a-c**), carbon utilization was perturbed for an extended period of time following sudden acetate addition. **a)** During the 150 min monitored following acetate addition (*t* = 0), OD_600_ nearly doubled, increasing exponentially at a rate of ∼0.3/h (red curve and right vertical axis). The concentrations of glucose remaining in the medium at various times, measured by HPLC (**Methods 3.3**), are shown as blue circles (left vertical axis). **b)** Glucose concentrations in panel **a** are plotted against OD_600_ as the blue circles. The best-fit linear fit of the data yields a specific glucose uptake of ∼11.6 mM/OD_600_ (negative of the slope of the blue line), giving a glucose yield of 0.09 OD_600_/mM glucose). [To further express the yield in term of biomass, one can use the conversion that 1 ml of OD_600_ culture corresponding to ∼500 mg of dry mass (*25*).] The corresponding data for steady state growth before and after acetate addition are plotted as the green and brown symbols, respectively. Thus, the specific glucose uptake in this period is more than double that of pre- and post-stress steady state. **c)** The rate of glucose uptake, obtained as the product of uptake per OD_600_ and the growth rate. The growth rate in the steady states before and after acetate addition was 0.93/h and 0.30/h, respectively. **d)** After the addition of acetate, cells began excreting pyruvate steadily, reaching almost 3 mM in the medium after ∼150 min. **e)** Alanine was also excreted, but at ∼100x smaller magnitude than pyruvate, reaching a maximum of ∼30 μM in the medium. The cells begin to take up the alanine after 120 minutes, suggesting the beginning of adaptation. **f)** Plotting the pyruvate concentration in the medium against OD_600_ at the corresponding time yields the pyruvate excretion per biomass, which was ∼16 mM/OD_600_ throughout this period (slope of the blue line). The corresponding data for pyruvate in steady state growth before and after acetate addition are shown as the green and brown symbols, respectively. Very little pyruvate was excreted in either steady state. **g)** Acetate in the medium was nearly unchanged during this period (blue circles, 0.2 mM in 150 min). **h)** The external acetate concentration in panel g is plotted against OD_600_ for the period following shift (blue circles). The slope gives the specific acetate uptake per OD_600_. The corresponding data for acetate in steady state growth before and after acetate addition are shown as the green and brown symbols, respectively. **i)** The rate of acetate uptake/excretion before, during, and after acetate addition. These rates are obtained by multiplying the specific acetate uptake/excretion by the corresponding growth rate. The specific acetate excretion before shift and the specific acetate uptake after shift are obtained from the slopes of the data in panel h.

**Figure S5.**
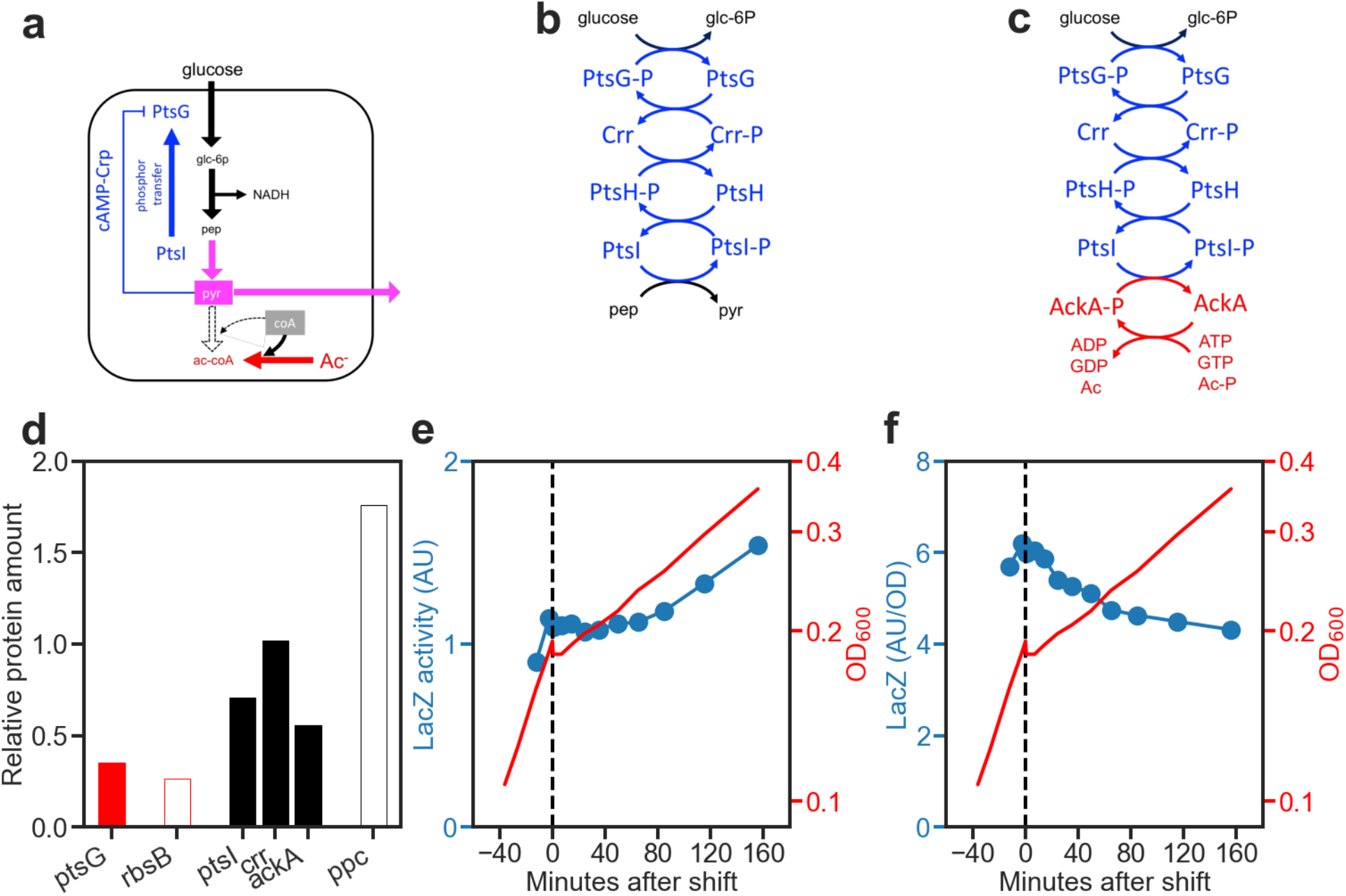
The slow adaptation in glucose uptake and pyruvate excretion. The heavy excretion of pyruvate (**Fig. S4d**) following acetate addition suggests that the cell experienced a bottleneck downstream of pyruvate. This likely resulted from the reduced rate of converting pyruvate to acetyl-CoA, due to the depletion of the CoA pool as large amounts of acetate entered the cytoplasm, leading to a buildup of pyruvate which subsequently leaked out of the cell (panel a). Somewhat surprisingly, the increase in internal pyruvate pool did not stop glucose uptake as expected of PTS-based glucose transport system (*67*), which couples the pep-to-pyr conversion to the transport and phosphorylation of glucose (blue arrows in panel a, with details in panel b): **Fig. S4c** shows a glucose uptake rate exceeding 3 mM/OD_600_/h in the 150 min following acetate addition, in between the uptake rate in the pre- and post-acetate steady state (∼4.5 mM/OD_600_/h and ∼1.5 mM/OD_600_/h, respectively). This modest reduction in glucose uptake despite a huge pyruvate buildup might be due to the existence of an alternative driver of PTS involving the enzyme Acetyl Kinase (AckA) (*68*) as illustrated in panel c. In this case, we expect the eventual reduction in glucose uptake (and the accompany stoppage of pyruvate excretion) in the post-acetate steady state to arise from the reduced level of the glucose transporter PtsG itself. Indeed, our proteomic data shows that the level of PtsG in the steady state with 30mM acetate dropped to ∼1/3 of that in the absence of acetate (panel d, filled red bar). Reduction in PtsG expression is expected to result from cAMP-Crp based transcriptional regulation which the promoter of *ptsG* is known to be controlled by (*69*); additionally, cAMP synthesis is known to be inhibited by the accumulation of keto-acids including pyruvate (*35*), thus linking the downregulation of PtsG expression to the increased pyruvate pool (blue inhibitory line in panel a). Indeed, other well-known member of the Crp-regulon such as RbsB (*70*) also exhibited substantially reduced levels in the post-acetate steady state (panel d, open red bar), while enzymes of glycolysis and the other PTS components changed little (filled black bars). We directly tested the reduced cAMP hypothesis by monitoring the expression of IPTG-induced LacZ following acetate addition. Synthesis of LacZ, quantified by beta-galactosidase activity (*35*), stopped for nearly an hour right after acetate addition before recovering (panel e). However, as the biomass (OD_600_) increased by only ∼20% in the first hour, the concentration of LacZ, proportional to LacZ amount per OD_600_ (since the cytoplasmic water volume per OD_600_ is nearly constant, (**Fig. S2e**)), decreased only moderately during this period, and continued to decline gradually afterward (blue circles, panel f). This requirement for “dilution” may contribute to why a long adaptation time exceeding the cell doubling time was needed for the reduction in glucose uptake and the eventual stoppage of pyruvate excretion. Another possible cause contributing to the slow reduction in pyruvate excretion is the demand for ATP, which is likely limited during the post-acetate transient due to the lack of pyr to ac-CoA conversion (**Fig. S4a**) and the lack of acetate uptake (blue bar, **Fig. S4i**). In this case, NADH produced by the additional glycolytic flux ending in the excreted pyruvate becomes an important source of energy. This need for NADH generation via pyruvate would be alleviated in the post-acetate steady state where acetate is taken up (brown bar, **Fig. S4i**) and TCA cycle can be used to generate energy. The restoration of TCA cycle flux requires the increase of Ppc level (**Fig. S6a**) which indeed occurs in the steady state with acetate (open black bar, panel d). As Ppc expression is negatively regulated by Cra (*71*) which is itself activated by cAMP-Crp (increasing under carbon limitation (*70*)), the increase of Ppc level may be slow due to the slow dilution of Cra when cAMP level dropped upon the addition of acetate (**Fig. 2h**).

**Figure S6.**
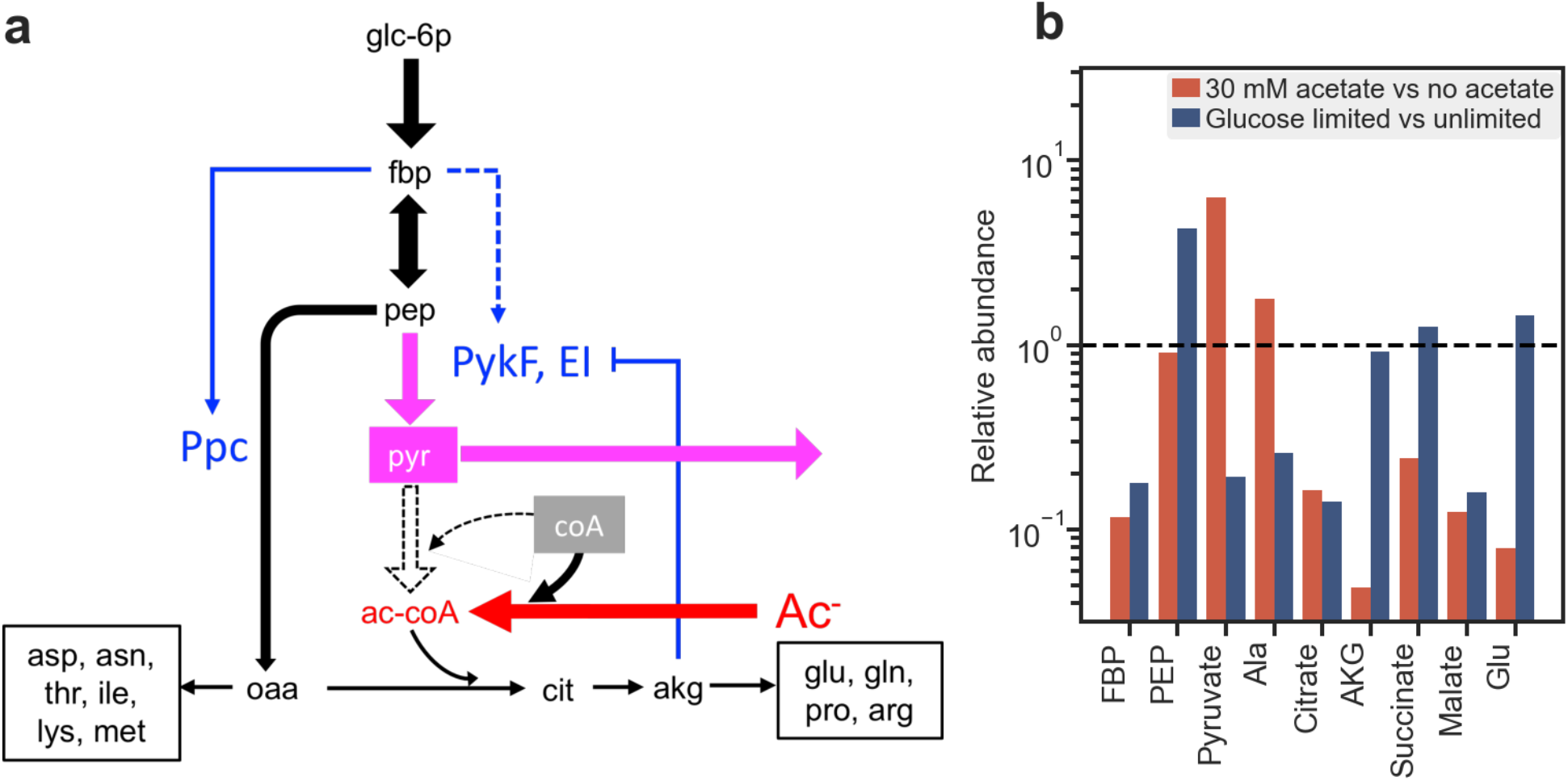
Reduction in anaplerotic flux immediately following acetate addition. The extended excretion of pyruvate (**Fig. S4d**) contrasts starkly with the very brief excretion of glutamate (**Fig. 2b**) following acetate addition. The latter is attributed to the strong reduction of the flux from pep to oaa (the anaplerotic flux) compared to the flux from pep to pyruvate as illustrated in panel a above. As described in **Fig. S5**, the pep-to-pyr flux cannot be converted to ac-CoA due to the lack of CoA and is therefore excreted out of the cell. Mechanistically, what favors the conversion of pep to pyr (catalyzed by Pyruvate Kinase, PykF, and by Enzyme I of PTS, PtsI) instead of pep to oaa (catalyzed by Phosphoenolpyruvate Carboxylase, Ppc) can be understood from the allosteric regulation of these enzymes, particularly, the rapid response of Ppc to fbp (*37*).

1. Both Ppc and PykF are activated by fbp (blue arrows above), but their sensitivity ranges are very different: K_A_ of Ppc for fbp is ∼20 mM (*36*). For fbp in the range of 20 mM to 5 mM (based on the absolute concentration in Ref. (*32*) and the relative change due to acetate addition shown in **Fig. 2d**, see also **Table S3a**), a drop in the fbp pool would significantly reduce the activity of Ppc; this has been directly characterized during glucose depletion by Xu et al (*37*). On the other hand, K_A_ of PykF is below 0.5 mM (*72*), making it insensitive to this range of changes in fbp concentrations. Furthermore, PtsI is not affected by fbp. Thus, change in fbp is not expected to affect the pep-to-pyr flux. We hypothesize that the fbp pool was quickly dropped in the first ∼10 min following acetate addition, from 20 mM down to the lower range, while the pep pool remained unchanged (similar to its steady state pools shown in **Fig. 2d**). As a result, the pep-to-pyr flux would hardly be affected but the pep-to-oaa flux would drop drastically (*37*). The latter would reduce the synthesis flux of all the TCA intermediates, as well as the 10 amino acids derived from them, as listed in the two boxes in panel a including glutamate. Thus, we expect these TCA intermediates and the derived amino acids to drop quickly to their steady state levels (**Fig. 2e,f**) in a few minutes after acetate addition, similar to those directly measured for the glutamate and glutamine pools (**Fig. 2c**).
2. The conversion of pep to pyr by PtsI is inhibited by akg (blue inhibitory line in panel a) (*73*). However, the drastic drop of TCA intermediates including akg disables this inhibition, reinforcing the channeling of flux from pep to pyr instead of oaa. This situation is to be contrasted to the response to a reduction in the external glucose supply. In this case, the fbp pool dropped similarly as under acetate stress according to our metabolomic data for glucose-limited growth (blue bars in panel b). But the pep pool increased 4-fold while the pyr pool decreased 5-fold. These changes are attributable to 1) the decrease of the glucose influx via the PTS system as the glucose-to-g6p conversion is reversibly connected to the pep-to-pyr conversion (**Fig. S5b**), and 2) the inhibition of PtsI by akg (*73*) (whose pool remains high; see below). The decreased pyr pool would account for the decreased alanine pool under glucose-limited growth, as the two pools are connected reversibly through transamination. The increased pep pool would not affect PykF, whose Michaelis constant for pep is only 0.03 mM according to Ref. (*72*) and 0.08 mM according to Ref. (*74*); nor would it affect PtsI, whose Michaelis constant is 0.18 mM (*75*). (The pep concentration in the unstressed condition was ∼0.2 mM; see **Table S3a**.) However, it is expected to increase the pep-to-oaa flux proportionally as the Michaelis constant of Ppc for pep is several mM (*36*); this would compensate for the loss of allosteric activation due to the drop of the fbp pool mentioned in the previous paragraph. The net result is the maintenance of the anaplerotic flux despite the reduced glucose influx, leading to the maintenance of the pools of the TCA intermediates including akg, and even an increased glutamate pool (blue bars, panel b). In summary, the metabolic responses to acetate stress and glucose limitation are opposite (panel b): Acetate stress resulted in the severe reduction of the pools of TCA intermediates and the amino acids derived from them, most notably glutamate, while keeping a very high pyruvate pool. In contrast, glucose limitation resulted in the reduction of the pyruvate pool and the amino acids linked to it, e.g., alanine, while maintaining the pools of TCA intermediates and particularly a high glutamate pool. The opposite response to these two perturbations is attributed primarily to the very different pep:pyr ratio (reduced by ∼6-fold under acetate stress and increased by ∼20-fold under glucose limitation). The pep:pyr ratio can in turn be traced to the different nature of the perturbations, an increase of the pyruvate pool due to the sequestration of CoA by acetate influx under acetate stress, and an increase of the pep pool due to the difficulty to transporting glucose under sugar uptake limitation (where we reduced the expression level of PtsG (*29*). The two opposite responses are reinforced by the inhibition of pep-to-pyr conversion by akg: a high pep-to-oaa flux maintains the akg pool which inhibits pep-to-pyr conversion, while a low pep-to-oaa flux depletes the akg pool and increases the pep-to-pyr flux.

**Figure S7.**
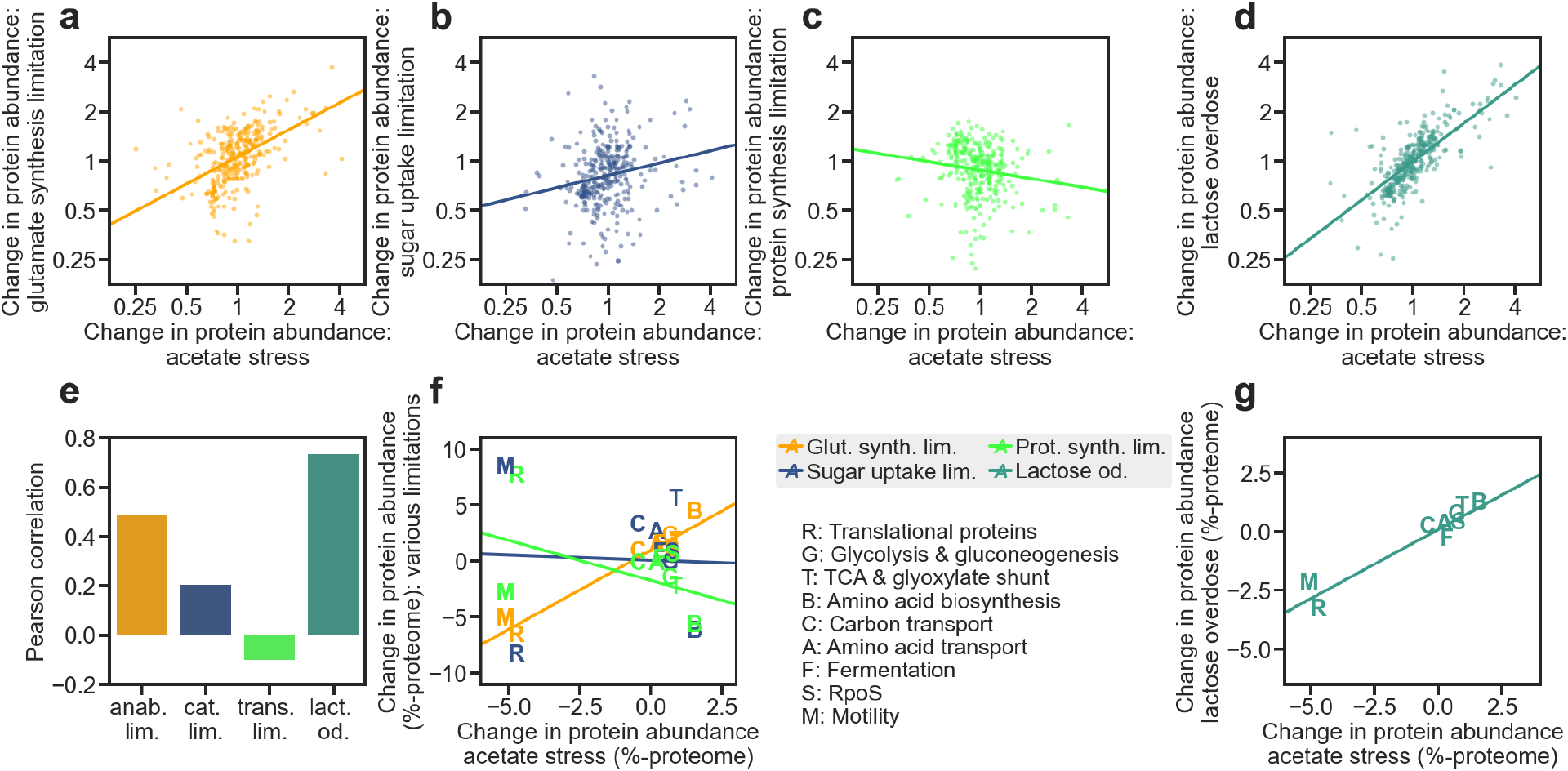
Proteome of acetate stressed cells. **a-d)** show scatter plots of fold-change in the abundance of each detected proteins between reference condition (glucose minimal medium, growth rate = 0.9/h), and a condition or strain of reduced growth at ∼0.4/h. x-axis: growth reduction due to acetate stress (30 mM sodium acetate at pH 6 in phosphate buffered medium); y-axis: growth reduction due to titratable expression of enzymes limiting glutamate synthesis (panel a), limiting sugar uptake (panel b), addition of Chloramphenicol providing sublethal inhibition of protein synthesis (panel c), lactose overdose (panel d). The proteome data for anabolic limitation, carbon limitation, and translational inhibition were obtained from Ref. (*29*). The proteomic data for acetate stress were generated for NCM3722 cells grown in phosphate buffered glucose medium (pH 6) supplemented with 30 mM sodium acetate; the data are shown in **Table S2a**. The proteomic data for lactose overdose (**Fig. S9**) were generated for HE620 cells grown in 25 ng/mL cTc with 750µM lactose added to the media; the data are shown in **Table S2b**. See **Methods 6**. **e)** Pearson correlation coefficient between the fold-changes in protein abundance due to acetate stress, and due to limitations in glutamate synthesis (orange), sugar uptake (blue), protein synthesis (green), and due to lactose overdose (teal) are shown in panels **a-d**. **f)** Scatter plots of the changes in the total abundances of proteins in each of the functional classes, between acetate stress and glutamate synthesis limitation (orange), sugar uptake limitation (blue), and translational inhibition (green). The functional classes are as described by the legend, e.g., “R” for translational proteins. The membership of proteins belonging to each class is based on Ref. (*70*) and are listed in **Table S2c**. The total abundances of detected proteins in each class under different growth limitations are shown in **figS. S8** and **Table S2a,b**.

**Figure S8.**
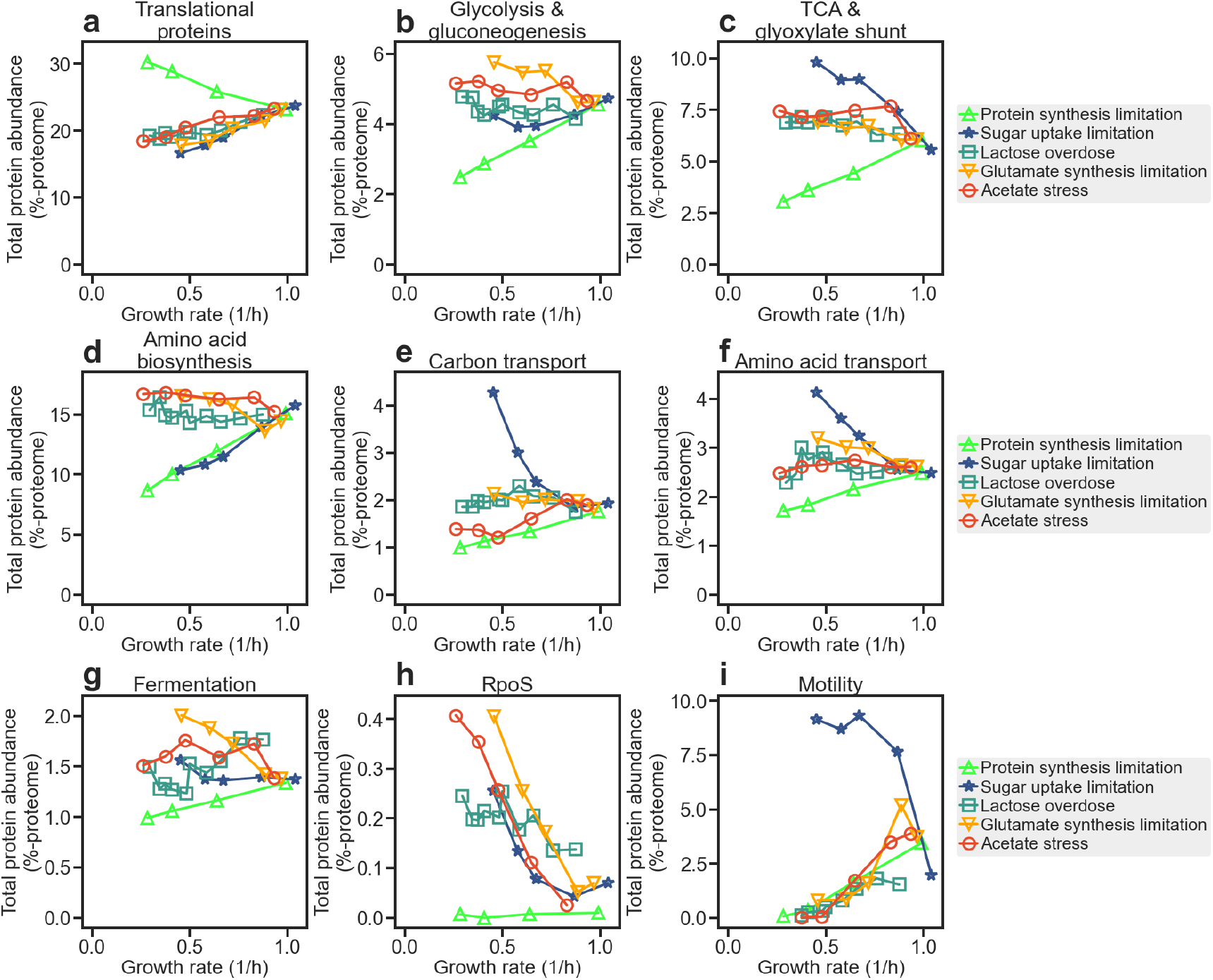
Functional grouping of proteome for acetate stressed cells. **a-i)** The proteome data shown in **Fig S7a-d** are grouped into a number of functional classes; see **Table S2c** and Ref. (*70*). The total abundance of proteins in each class, measured as the fraction of total protein mass, is plotted against the growth rate for each type of growth limitation: acetate stress (red circles), sugar uptake limitation (blue stars), glutamate synthesis limitation (orange down-triangle), translational inhibition (green up-triangles), lactose overdose (teal squares). The proteome data for glutamate synthesis limitation, sugar uptake limitation, and translational inhibition were obtained from (*29*). The proteomic data for acetate stress were generated for NCM3722 cells grown in phosphate buffered glucose medium (pH 6) supplemented with different concentrations of sodium acetate; the data are shown in **Table S2a**. The proteomic data for lactose overdose were generated for HE620 cells grown in 25 ng/mL cTc with various concentrations of lactose added to the media; the data are shown in **Table S2b**. See **Methods 6**.

**Figure S9.**
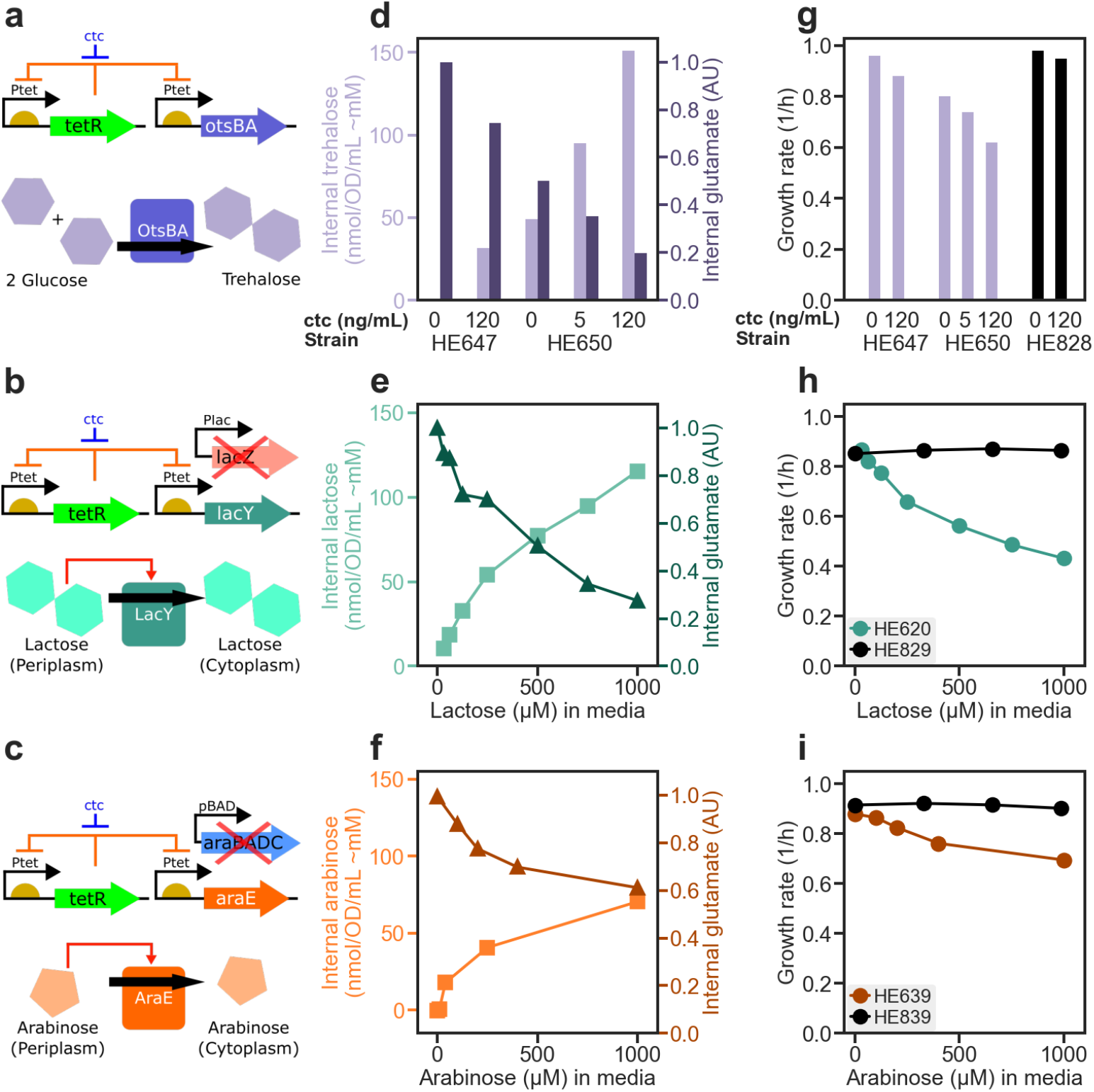
Overdose of useless metabolites. **a)** Schematic of the construct for the titratable accumulation of internal trehalose. Strains HE647 and HE650 were constructed to have titratable expression of the *otsBA* operon, which encodes trehalose-6-phosphate synthase and trehalose-6-phosphate phosphatase (*76*) driven by the P_Ltet_ promoter (*52*). P_Ltet_ was controlled by TetR expression, which was provided by 3 copies of *tetR* driven by P_Ltet_ on the chromosome. The two strains differ in that *otsBA* is either expressed from the chromosome (HE647) or from a plasmid pZA31 (HE650). This expression system is capable of driving protein output by over 100x in response to changes in the level of an inducer, chloro-tetracycline, added to the media (*35*). **b)** We similarly constructed strain HE620 which allows titratable accumulation of lactose: A plasmid harboring titratable expression of *lacY* (encoding the Lac Permease, LacY) by the P_Ltet_ promoter is transformed into cells containing one copy of *tetR* driven by P_Ltet_, which are themselves derived from NCM3722 with a chromosomally inserted *tetR* driven by the P_Ltet_ promoter, but with additionally the deletion of chromosomal *lacIZY*. Since they lack LacZ, HE620 cells in media with lactose will import lactose but not degrade it, resulting in lactose accumulation. **c)** The arabinose overdose strain (HE639) was designed similarly to the lactose overdose strain. A plasmid harboring titratable expression of *araE* (encoding the Ara Permease, AraE) by the P_Ltet_ promotor is transformed into cells containing one copy of *tetR* driven by P_Ltet_. Additionally, the *araBAD* and *araC* genes were removed from the chromosome to avoid arabinose metabolism. **d)** The accumulation of trehalose (light purple bars, left y-axis) in trehalose titration strains (HE647 and HE650) grown in glucose minimal medium fixed to pH 6, with various inducer (cTc) concentrations and no acetate. The internal glutamate pool in these cells as measured by HPLC are shown as dark purple bars (right y-axis). **e)** The accumulation of lactose (triangles, left y-axis) by the lactose titration strain HE620 grown in phosphate buffered glucose media with 25 ng/mL ctc to turn on LacY. The concentration of lactose supplemented in the medium is shown as the x-axis. The corresponding internal glutamate concentrations are shown as squares (right y-axis). **f)** The accumulation of arabinose (squares, left y-axis) by the arabinose titration strain HE639 grown in phosphate buffered glucose media. The concentration of arabinose supplement in the medium is shown as the x-axis. The corresponding internal glutamate concentrations are shown as triangles (right y-axis). **g)** Growth rates of the trehalose titration strains grown in conditions described in panel c are shown as the dark purple bars. The growth rates of the control strain HE828 (identical to HE650 except that *otsBA* is replaced by *gfp*) are shown as black bars. **h)** Growth rates of the lactose titration strain HE620 grown in conditions described in panel e are shown as circles. The growth rates of the control strain HE829 are shown as black circles. HE829 is constructed identically to HE620 except the *lacY* on the plasmid was replaced with *gfp*. **i)** Growth rates of the arabinose titration strain HE639 grown in conditions described in panel f are shown as circles. The growth rates of the control strain HE839 are shown as black circles. HE839 is constructed identically to HE639 except the *araE* on the plasmid was replaced with *gfp*.

**Figure S10.**
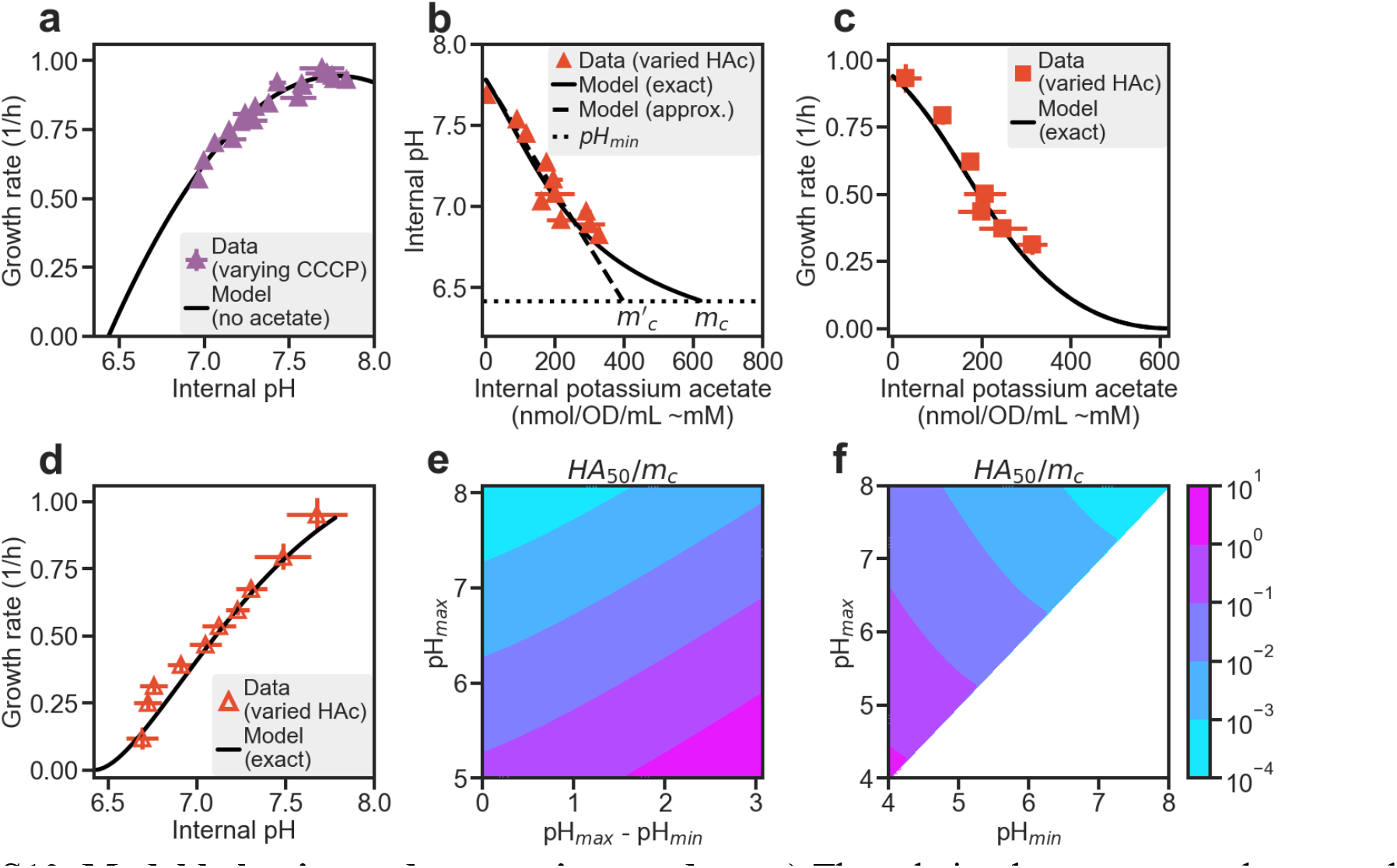
Model behavior and comparison to data. **a)** The relation between growth rate and internal pH, as revealed by the CCCP experiment in the absence of acetate (same data as the purple triangles in **Fig. 3e**), is well described by the quadratic form (solid line) given by Eq. (4), with the best-fit parameters pH_max_ = 7.78, pH_min_ = 6.42, and λ_0_ = 0.94 h^-1^. **b)** According to the solution of the model (**Supp Note**), the relation between internal pH and potassium acetate concentration is expected to be approximately linear for internal pH spanning as much as half of the range between the normal value pH_0_ and the minimal value pH_min_; see the dashed line, which extrapolates to an apparent inhibition concentration 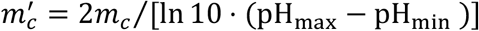 where internal pH reaches its minimal value and growth rate vanishes. Our data is best fitted by *m_c_* ≈ 620 *mM* (corresponding to 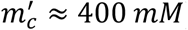). This fixes the lone unknown parameter of the model. Using this parameter value, the exact solution of the model is shown as the solid line. **c)** The model with the fixed parameters accurately describes the relation between internal potassium acetate and growth rate. **d)** The model with the fixed parameters accurately describes the relation between internal pH and growth rate due to acetate stress. **e)** and **f)** provide two plots of the dependence of the half-inhibitory acetic acid concentration (*HA*_50_), defined as the concentration of acetic acid (external or internal) where the growth rate is reduced to 50% of the unstressed value. The half-inhibitory concentration is plotted for different values of the cellular pH in stress-free conditions (pH_max_) and minimal pH value (pH_min_), and the result is expressed relative to a characteristic internal potassium acetate concentration, *m_c_*. Data in panels a, b, c, and d are grouped according to the same applied stress level and contain n ≥ 3 biological replicates. Error bars are calculated from the standard deviation.

### Supplementary Tables

**Table S1a:**
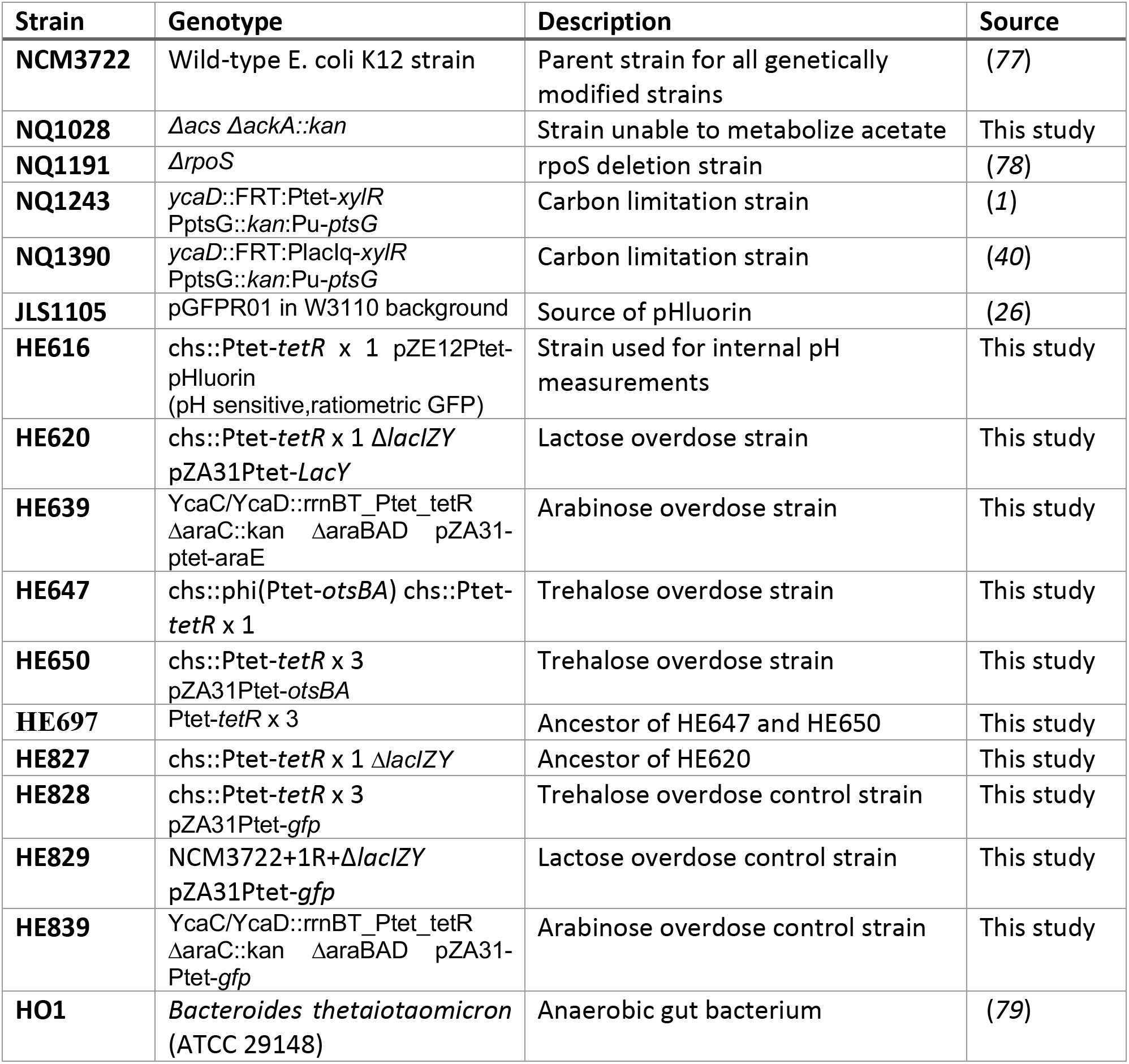
Strain used in this study.

**Table S1b.**
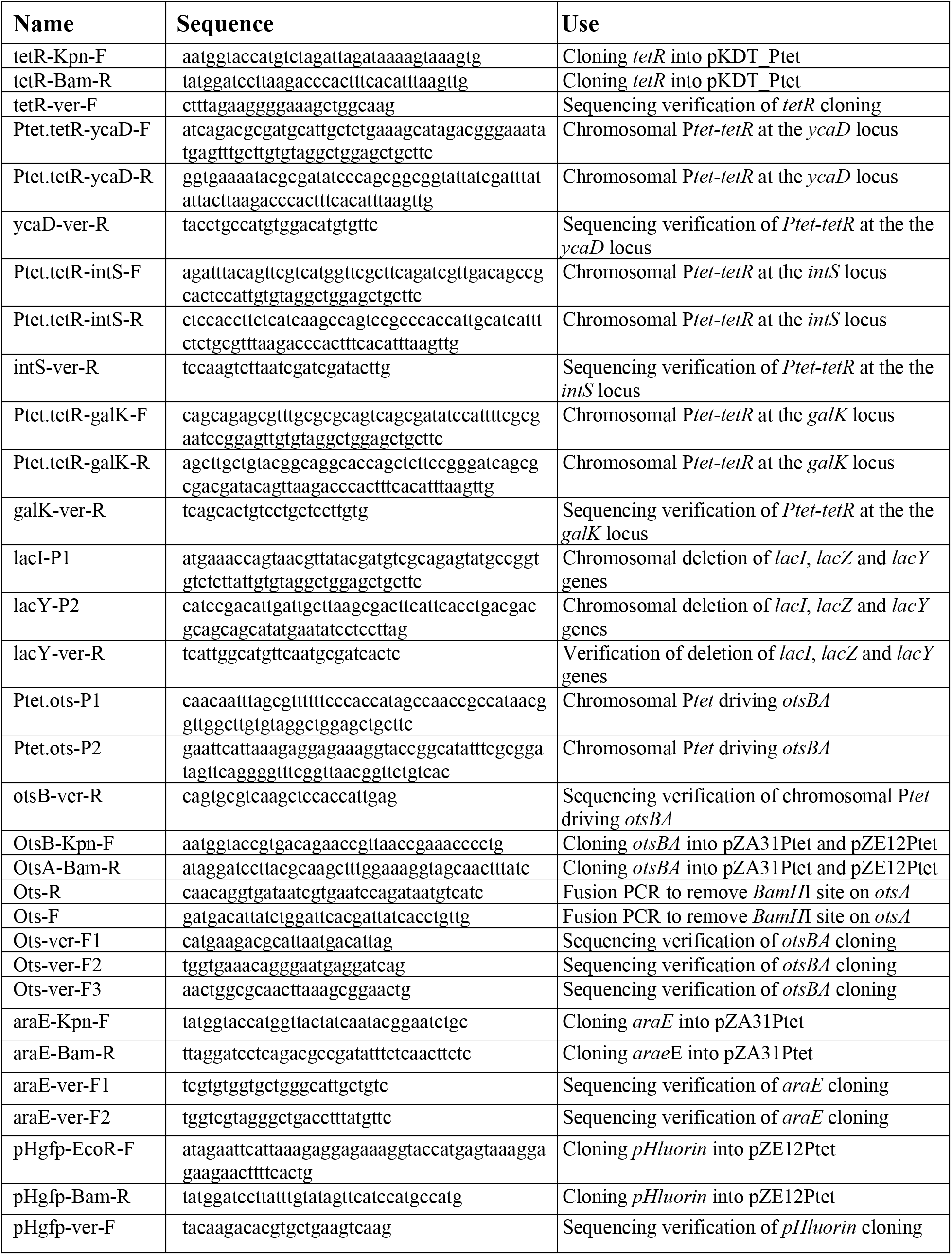
Oligonucleotides used in this study.

**Table S2a.** Proteomic data for acetate stress. Relative and absolute proteomic data for NCM3722 cells where growth rate is limited by the addition of different concentrations of sodium acetate.

**Table S2b.** Proteomic data for lactose overdose. Relative and absolute proteomic data for HE620 cells where growth rate is limited by the accumulation of lactose.

**Table S2c.** Membership of proteins belonging to different functional classes. Functional categories of proteins in the *E. coli* proteome used in **Fig. S7f,g** and **Fig. S8** based on (*70*).

**Table S3a.** Metabolomic data for NCM3722 cells under a range of acetate stress. Relative and absolute proteomic data for NCM3722 cells where growth rate is limited by the addition of different concentrations of sodium acetate.

**Table S3b.** Metabolomic data for NCM3772-derived cells under carbon-limited growth. Relative and absolute metabolite measurements for cells with growth limited by controlling carbon uptake through PtsG. Strains NQ1243 and NQ1390 were grown in phosphate buffered media. Expression of PtsG was controlled by addition of 3MBA.

**Table S3c.** Metabolomic data for NCM3772-derived cells under trehalose-overdose growth. Relative and absolute metabolite measurements for cells with growth limited by accumulation of trehalose by OtsAB.

